# Bioengineered human tonsil organoids as an immuno-engineering platform for evaluating immune functions

**DOI:** 10.1101/2025.10.04.680210

**Authors:** Intan Rosalina Suhito, Donovan Eu Kum Chuen, Hong Sheng Cheng, Nguan Soon Tan, Zhe Zhang Ryan Lew, Kai Sen Tan, Justin Jang Hann Chu, Giselle G.K. Ng, Yvonne C.F. Su, Gavin J.D. Smith, Chee Wah Tan, Andy Tay

## Abstract

Although animal models remain the preclinical gold standard, they are constrained by interspecies differences in adaptive immunity, including antibody class-switching, affinity maturation, and multicellular interactions. To address this gap, we established a more physiologically relevant platform, a bioengineered tonsil organoid (BTO) system that combines autologous tonsil-derived stromal cells with immune cells to generate highly uniform organoids, preserving multicellular complexity and sustained stromal-immune interactions. BTO cultures mounted robust, antigen-specific recall responses to influenza, tetanus, and COVID-19 antigens. Notably, stimulation with ovalbumin and SARS-CoV-1 spike protein expanded antigen-recognizing B cells, indicating capacity for naïve responses. Digital spatial transcriptomics and proteomics analysis confirmed immune-stromal crosstalk underpinning both humoral and cellular responses of BTO after challenge. These data highlight stromal cells as active scaffolds that not only maintain 3D architecture but also facilitate immune activation, memory recall, and recognition of naïve antigens. Together, BTO offers a human-relevant model for mechanistic immunology and for pre-clinical evaluation on immunotherapies and vaccines not achievable with 2D cultures or animal models.

## Introduction

Infectious diseases pose global-scale threats to human health, driven by a growing number of epidemic and endemic pathogenic viruses. The COVID-19 pandemic, which has killed more than 7.1 million people worldwide, underscores the urgency of high-throughput preclinical models that can rapidly characterise pathogens and accelerate discovery of effective therapeutics, including vaccines and anti-viral antibodies.^1^ Vaccination remains a cornerstone of public health strategy, priming protective immunity against current and future infectious agents.^2^ Yet. traditional vaccine development relies heavily on animal models despite interspecies differences in antibody class switching, affinity maturation, and somatic hypermutation that diverge from human adaptive immunity.^3^ These drawbacks contribute to clinical-stage failures when candidate vaccines are tested in humans.^4^

The FDA Modernization Act 2.0, enacted in 2022, promotes the use of advanced non-animal models (e.g., organoids or micro-physiological systems) for drug development and testing, which underscores the growing recognition of human-relevant platforms as powerful tools to study immune responses and therapeutic efficacy.^5^ *Ex vivo* 3D explant tissue culture preserve native organ architecture and can inform potential vaccine responses, but they are short-lived (∼3 days) due to the inability to sustain viable immune cells long-term.^6,7^ Access to healthy human tissues is also limited, as most samples derive from biopsy, surgical resections, or deceased donors. Explants are also non-renewable and cannot be propagated, hindering repeated experimentation on the same donor. A more physiologically relevant model, renewable model for translational human immunological studies is therefore needed.

Organoids are self-organizing multicellular 3D structures that recapitulate key structural and functional features of their tissue of origin.^8^ They can offer greater physiological relevance, longevity, scalability, and cellular complexity than 2D cell culture and tissue explants. Recent immune organoid advances, particularly from lymphoid tissues, enable human-centric studies prior to clinical translation.^9^ Tonsils, a readily available mucosal-associated lymphoid tissue from routine tonsillectomies, are especially attractive for modeling immunity.^10^ They are rich in B cells, T cells, dendritic cells, and stromal cells that support germinal center-like organization.^11^

Only a handful of human tonsil-derived immune organoids have been reported. Wagar *et al*. described the first human tonsil organoid culture but lacked stromal cells and matrix component, yielding loose aggregate of cells that complicate sectioning and high-content imaging.^12,13^ Standard well-insert techniques can also introduce variability in organoid size and morphology, leading to non-uniform drug diffusion, altered immune cell dynamics, and reduced experimental reproducibility.^14,15^ Kim *et al*. reported a human tonsil organoid with innate immune responses to pathogen-derived molecules, but bias differentiation toward epithelium limited evidence for human adaptive immunity.^16^ Most recently, Zhong *et al.* developed synthetic immune organoids derived from healthy donors and patients with B cell lymphoma using a biomimetic hydrogel system.^17^ However, the matrix degraded by more than 50% within four days, and incorporation of non-autologous CD40L-expressing stromal cells and human tonsil-derived follicular dendritic cells (HK-FDCs) may compromise authenticity and HLA compatibility.

Stromal cells are essential organizers of lymphoid tissue architecture, guiding immune cell migration, activation, and retention.^18^Among them, fibroblastic reticular cells (FRCs) form conduit networks that support antigen transport and cytokine gradients.^19,20^ Studies have shown that in the absence of FRCs, both humoral and cellular immunity toward viral evasion are compromised.^21,22^ FRC leads to declined immune responses because of stromal architecture and functional disruption, which affect downstream processes like immune cell activation, antigen delivery, and immune surveillance. ^23,24^ Despite their importance, FRCs remain underexplored in immune organoids. We therefore hypothesize that incorporating autologous stromal cells would yield a more physiologically authentic human immunity model with sustainable matrix, high multicellular diversity, preserve immune memory, and the capacity to recognize naïve antigens.

In this study, we establish a bioengineered tonsil organoid (BTO) by incorporating and optimizing ratio of autologous tonsil stromal cells (TSCs) and tonsil-derived cell mixtures (TCMs). Using a hanging drop method, we generated uniform organoids with multicellular diversity comparable to *ex vivo* tonsil tissue. Fibroblastic reticular cells (FRCs) provided structural support and primed immune cells to deploy immune responses against antigenic challenges. BTOs recapitulated recall humoral responses to SARS-CoV-2 (COVID-19), tetanus, and influenza A virus, and mount naïve responses to ovalbumin and SARS-CoV-1 spike protein, including production of antigen-specific neutralizing antibodies. Digital spatial transcriptomic and proteomic analyses revealed immune-stromal crosstalk underpinning these responses and sustaining immune surveillance. Together, BTO advances physiological fidelity over existing organoid models and provides a practical platform for human immunology and preclinical screening of immunotherapeutics.

## Results

### Bioengineered tonsil organoids (BTOs) can be reproducibly established

We developed a bioengineered tonsil organoid (BTO) model by optimizing the ratio of autologous tonsil stromal cells (TSCs) within tonsil-derived cell mixtures (TCMs). Tonsil tissue was enzymatically digested to obtain single cell suspension named as TCMs (donor details in **Table S1**). TSCs were subsequently isolated by plating TCMs, removing lymphocytes, and expanding adherent stromal cells (**Fig 1a**).

**Figure 1.**
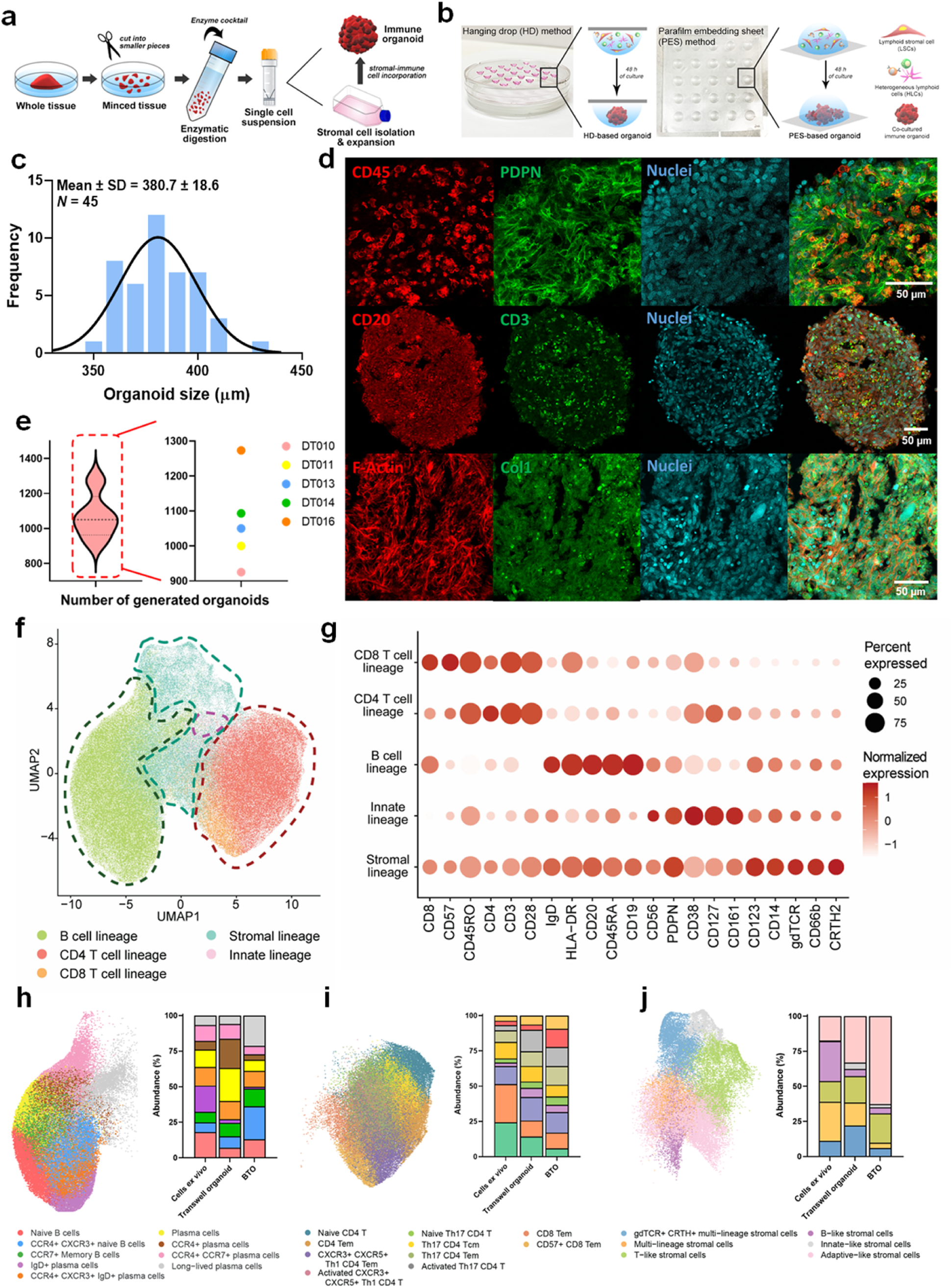
Addition of autologous stromal cells create morphologically uniform bioengineered tonsil organoids. Schematic illustrating the workflow for **(a)** tonsil tissue processing for FRCs isolation and organoid culture and **(b)** tonsil organoid generation with hanging drop (HD) and parafilm embedding sheet (PES) methods. **(c)** Bar graph showing size distribution of HD-based tonsil organoids (380.7 ± 18.6 µm) with a 10:1 ratio (n=45). **(d)** Immunofluorescence images of sliced organoid showing cell-to-cell interactions between TSCs and TCMs, with immune cells distributed across the stromal microenvironment that is rich with endogenously produced collagen. Red: PDPN expression corresponds to TSCs. Green: CD45 expression corresponds to immune cells in TCMs. **(e)** Number of generated organoids per ± 1 cm^3^ of tonsil tissue from five donors, with an average of Mean ± SD. **(f)** UMAP data of cell populations in HD tonsil organoids compared to that in native tissue and transwell organoids. **(g)** Dot plot of differentially expressed markers for immune- and stromal cell subsets in all sample groups, indicating preservation of all major immune cell types including naïve B and T cells, B and T cell memory, helper T cells, and dendritic cells. UMAP data showing clusters and relative abundance percentage of multiple cell subsets corresponding to **(h)** B cell lineage, **(i)** T cell lineage, and **(j)** stromal cell lineage in the three groups. Data in **(f)** to **(j)** are based on cytometry by time of flight (CyTOF) analysis.

Immunofluorescence staining confirmed TSC morphology with podoplanin (PDPN) and cytoskeletal protein F-actin expression, consistent with a FRC phenotype (**Fig S1a**). The TSCs underwent further characterizations by flow cytometry, confirming their identity through high expression of stromal mesenchymal markers such as CD73, CD90, CD105, and PDPN, and low expression towards hematopoietic markers (CD14, CD34, CD45), epithelial marker (ECAD), and endothelial markers (CD31) as shown in **Fig S1b**. We also compared the hanging drop (HD) and parafilm embedding sheet (PES) methods for spontaneous organoid formation across multiple ratios of TCMs and TSCs (**Fig 1b** (**Fig S1c-e**). Based on our results, the HD method with 10:1 ratio yielded optimal organoid dimensions by maintaining TSC viability and growth, thereby, providing structural integrity and support for immune cells.

The HD approach produced uniformly spherical organoids that enabled simple and reliable readouts. This morphology is consistent with reported brain, heart, lung and kidney organoid models created with matrix materials such as Matrigel.^25–28^ It offered a technical advantage over transwell aggregates or PES culture, which formed heterogeneous and irregular organoids. Size distribution analysis indicated consistency in BTO formation using HD method, which is a crucial factor for reproducible downstream characterization assays (**Fig 1c**). Immunofluorescence staining revealed spatial arrangement and stromal-immune cells interactions. The expressions of CD45^+^, CD20^+^, and CD3^+^ clearly indicated the even distribution of immune cell subsets within the stromal-rich organoid matrix. Additionally, co-expression of F-actin and collagen type I (Col1) illustrated the structural integrity and extracellular matrix (ECM) provided by stromal cells (PDPN^+^) to facilitate essential cellular interactions that mimic native lymphoid tissue architecture (**Fig 1e**). Robust performance across multiple donors supports the generalizability of this protocol for human immunological studies and applications (**Fig 1d**).

### BTO model preserves multicellular diversity

To evaluate the multicellular composition of the BTO model, and to benchmark its fidelity against donor-matched native tonsil tissue and the first reported transwell tonsil organoid model,^12,13^ we performed high-dimensional immune profiling using cytometry by time-of-flight (CyTOF). This allowed for comprehensive phenotyping of immune and stromal populations within our organoid system, revealing the extent to which cellular diversity and organization are preserved *in vitro*. The UMAP projection identified five major lineages, namely B cells, CD4⁺ T cells, CD8⁺ T cells, innate immune cells, and stromal cells (**Fig 1f**). Distinct clustering of these populations described clear lineage segregation and preservation of immune and stromal architecture. The dot plot in **Fig 1g** further analyzed lineage-specific marker expressions of those major cell types with row representing cell lineage, while column correspond to gold-standard surface markers. Notably, strong CD19 and CD20 expressions confirmed the presence of B cells. CD4, CD8, and CD45RO expressions indicated that T cell populations were present, while stromal cells were marked by high PDPN and CD90 expression, supporting the multicellular diversity of the BTO model.

Sub-classification UMAP of B cells, stromal cells, and CD4⁺ T cells provided a higher-resolution UMAPs that indicated heterogeneity within major cell types (**Fig 1h-j**). In **Fig 1h**, UMAP analysis of B lineage cells revealed a spectrum of phenotypes, including naive B cells, memory B cells, plasma cell subsets, and long-lived plasma cells, which captured native tonsillar B cell diversity. The side-by-side bar chart showed quantification of relative abundance of these subsets across three different groups, including tonsil cells from freshly digested *ex vivo* tissue, transwell cultured organoid, and BTO. Compared to the transwell technique, the BTO maintained a more balanced composition of naive and memory B cells, as well as long-lived plasma cells, suggesting improved support for B cell differentiation. **Fig 1i** shows the diversity of T cell populations by identifying multiple functionally relevant subtypes such as naive T cells, regulatory T cells, and activated effector memory subsets. Relative proportions of all T cell subsets across culture conditions showed that the BTO can support a broader range of T helper differentiation pathways comparable to *ex vivo* tissue. Lastly, the result in **Fig 1j** focused on stromal cell sub-populations, where distinct subsets including gdTCR⁺ CRTH⁺ multi-lineage stromal cells, B-like stromal cells, and T-like stromal cells were identified. The BTO preserved a greater number of stromal cell subsets compared to transwell culture, particularly dominated with adaptive-like stromal cells, indicating the inclusion of autologous FRCs in our system is critical for recapitulating the native tissue microenvironment.

Collectively, BTO models maintain rich multicellular diversity across B cells, T cells, and stromal cells, closely mirroring the cellular complexity of native tonsil organs. While protocol-dependent differences are expected (transwell versus BTO), we consistently observed superior preservation of naïve and memory B cells, long-lived plasma cells, and adaptive-like stromal subsets. The T cell sub-populations were largely comparable to *ex vivo* tonsil tissue, with a modest shift towards CD8^+^ over CD4^+^ T cells. Most importantly, explicit characterization of FRCs highlights stromal-immune interactions as a key determinant of physiological fidelity in lymphoid organoids.

### BTOs are activated by immunostimulatory cues

We evaluated the immunological functions of BTO by stimulating it with canonical mitogens and innate agonists (**Fig 2a**). Phytohemagglutinin (PHA) is a potent T-cell mitogen that crosslinks surface receptors to drive activation and proliferation.^29^ Upon PHA treatment, BTOs increased in size and compactness, as shown in the microscopic images and quantified in violin plot in **Fig 2b-c**. We also exposed BTOs to lipopolysaccharide (LPS), a Toll-like receptor 4 (TLR4) agonist that initiates innate immune cascades.^30^ Flow cytometry showed an increased survival of monocytes after LPS, with TLR4 recognition on monocytes that promote inflammatory cascade (**Fig S2**). Altogether, these data showed increased viability of immune cells within the 3D organoid confinement and were responsive to external cues. The controlled size and shape of BTOs further facilitate visualization of morphological responses to immunological stimuli.

**Figure 2.**
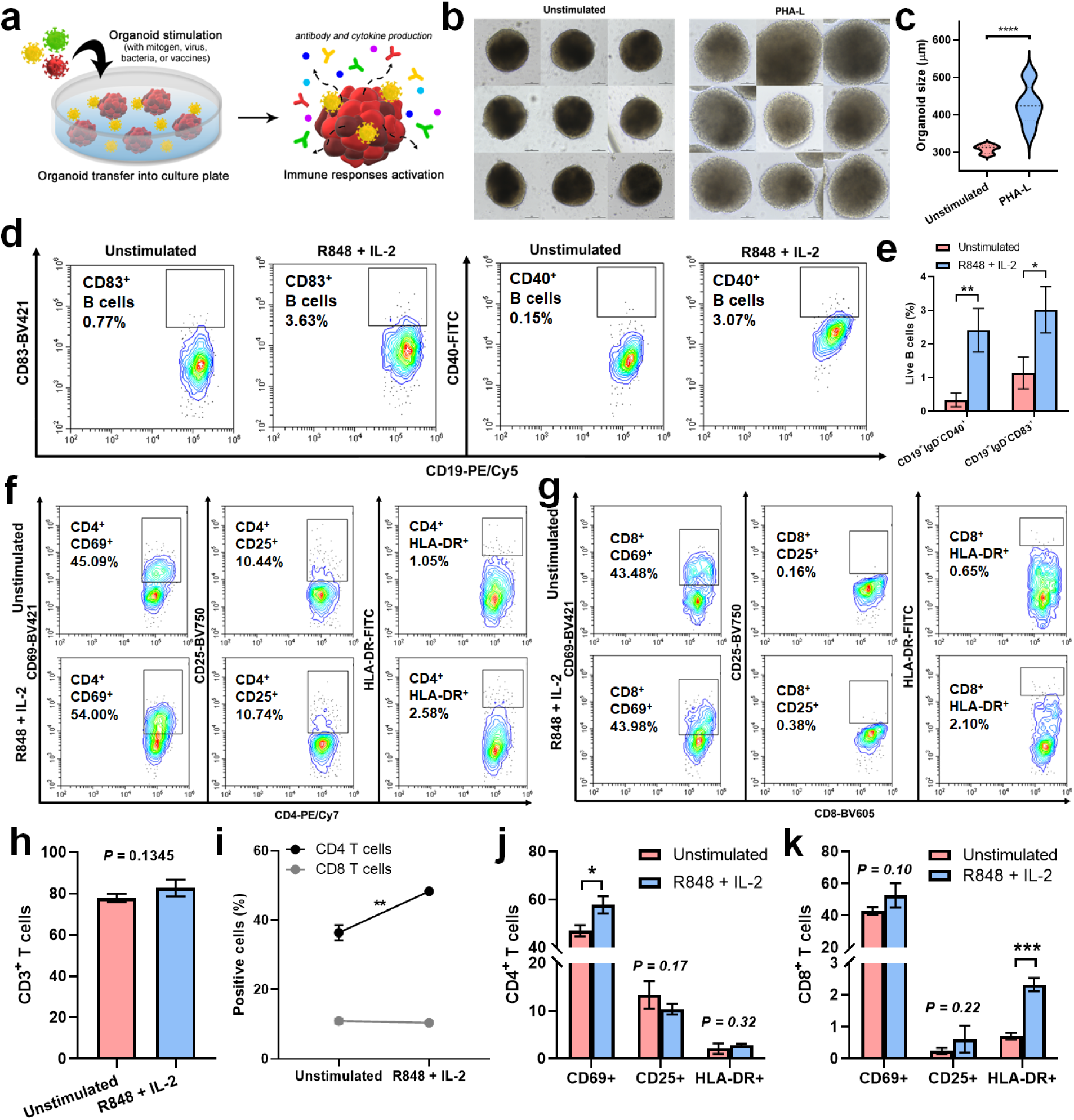
Immune cells in BTO respond to gold-standard agonist and cytokine stimulations. **(a)** Schematic illustrating tonsil organoid stimulation with molecules of interest to induce immune responses and activation. **(b)** Microscopic images (scale bar = 100 µm) and **(c)** violin plot showing proliferative capability of tonsil organoid after PHA stimulation. The controllable morphologies of HD organoids can enable simple readout based on size and shape changes. **(d)** Flow cytometry graphs and **(e)** bar graph indicating B cell activation in BTO after stimulation with R848 (1 µg/ml) and IL-2 (10 U/ml) cocktail. **(f-g)** Flow cytometry graphs of CD4^+^ T cells and CD8^+^ T cells, **(h)** bar graphs of total percentage of CD3^+^ T cells, **(i)** line graph showing changes in percentage of CD4^+^ and CD8^+^ T cells, and **(j-k)** T cell activation marker profiles from CD4^+^ and CD8^+^ T cells in tonsil organoid after stimulation with immunostimulatory cocktail consisting of R848 (1 µg/ml) and IL-2 (10 U/ml). All data are presented as mean ± S.D. Unpaired, two-tailed t-test was performed to calculate statistical significance. \**P* < 0.05; ** *P* < 0.01; \*\*\**P* < 0.001; **** *P* < 0.00001; no significance (n.s.), *P* > 0.05.

The BTOs were further tested for their ability to undergo immune activation following stimulation with TLR7/8 agonist (R848) and IL-2. Flow cytometry revealed clear B-cell activation within the organoid (**Fig 2d-e**). Specifically, CD19⁺ B cells upregulated CD40 and CD83 markers associated with germinal-center co-stimulation and B cell activation through B cell receptor (BCR), TLRs or CD40 engagement.^31,32^ These data show that BTOs preserve functional B cell signaling and provide a tractable system for studying B cell biology *in vitro*. This B cell response also underscored the physiological relevance of the organoids as R848 mimics viral RNA sensing through innate pathways while IL-2 provides essential T cell-derived signaling.

In addition to B cells, the T cell compartment also exhibited robust activation (**Fig 2f-k**). A modest increase in CD3⁺ cell numbers after stimulation (**Fig 2h**) aligns with reports that higher IL-2 doses (20-500 IU/ml) are typically required for significant T cell expansion^33^ Relative proportions of CD4⁺ and CD8⁺ subsets shifted (**Fig 2i**), suggesting potential remodeling with tendency towards CD4^+^ T cell expansion. Activation markers were differentially upregulated (**Fig 2j-k**), including CD25 as IL-2 α chain receptor and HLA-DR as a marker of sustained activation and antigen presentation.^34,35^ The immunostimulatory cocktail seemed to activate CD8^+^ T cells earlier as reflected in T cell late activation marker (HLA-DR expression), while CD4^+^ T cells showed latency in cellular activation as only early activation marker (CD25 expression) was markedly increased. The data demonstrate that B and T cells within the BTO remain viable and responsive, mounting dynamic, stimulus-appropriate activation to gold-standard immunostimulatory molecules.

### BTOs retain donor immune memory

Given widespread vaccination and/or infection during the COVID-19 pandemic in Singapore, SARS-CoV-2 antigens are ideal for testing humoral recall responses. The receptor-binding domain (RBD) of spike protein is the principal ACE2-binding region and a key target of neutralizing antibodies.^36^ BTO stimulated with COVID-19 RBD protein elicited a modest increase in total B cells, with expansion of GC plasmablasts and plasma cell differentiation, demonstrating that BTOs were able to support antigen-driven B cell differentiation (**Fig 3a-b**, **Fig S3a-c**). The presence of GC plasmablast is particularly important as it marks a functional immune cascade from antigen recognition through GC activation to antibody-secreting cell development.^37,38^ Interestingly, RBD protein stimulation also triggered cellular immunity with activation of CD4⁺ and CD8⁺ T cells (**Fig S3d-h**).

**Figure 3.**
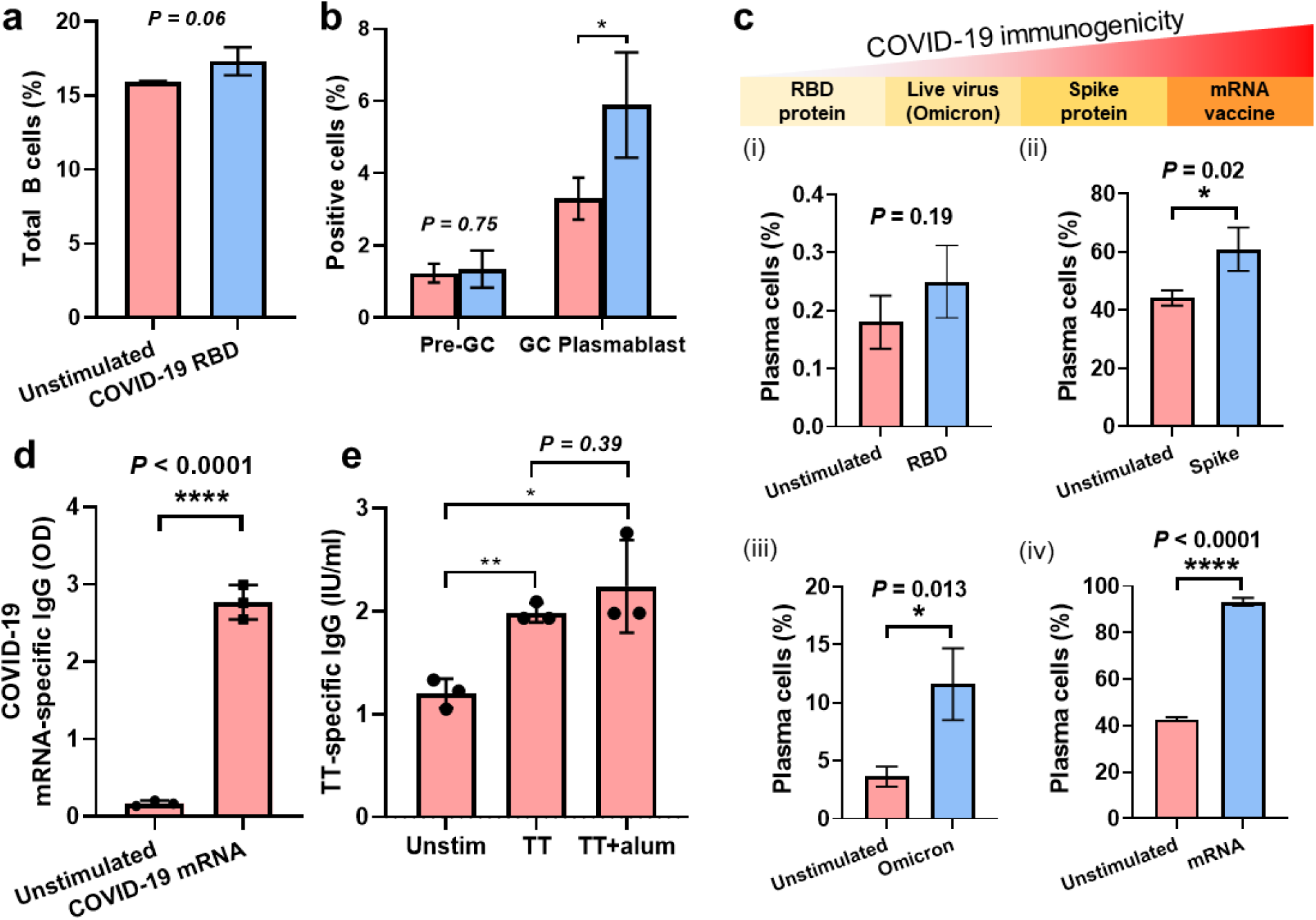
Tonsil organoids mount effective memory recall responses to previously exposed antigens. Quantification of **(a)** total B cells and **(b)** GC B cell responses in BTOs after COVID-19 RBD protein stimulation. **(c)** COVID-19 immunogenicity level from (i) RBD, (ii) spike protein, (iii) live virus (var. Omicron), and (iv) mRNA vaccine to direct plasma cell differentiation. COVID-19 mRNA antigen showed the highest potential to induce humoral immune responses. **(d)** ELISA data demonstrated a significant increase in COVID-19 mRNA-specific IgG production after stimulation. **(e)** ELISA data of tonsil organoid stimulation with tetanus toxoid (TT) protein, showing an increase in tetanus-specific IgG. All data are presented as mean ± S.D. Unpaired, two-tailed t-test was performed to calculate statistical significance. \**P* < 0.05; ** *P* < 0.01; no significance (n.s.), *P* > 0.05.

We next compared spike protein, live virus (var. Omicron), and an mRNA vaccine (Pfizer). A clear hierarchy of immunogenicity emerged (**Fig 3c**). The full-length spike protein elicited stronger B cell differentiation than RBD, while the COVID-19 mRNA vaccine produced the most robust plasma cell response, consistent with the capacity of lipid nanoparticle-encoded antigens to amplify humoral immunity through persistent GC B cells attributed to affinity maturation and breadth beyond convalescent infection.^39^ This pattern indicated clear differences in immunogenic potential between subunit proteins, whole live virus, and mRNA vaccines, suggesting that the BTO model faithfully reflects antigen-specific immune hierarchies. Antigen-specific ELISA corroborated these findings. ELISA quantification data found significant increases in COVID-19 mRNA-specific IgG production after stimulation (*p*<0.0001), demonstrating B cell functions in BTO to generate protective antibody upon memory recall response (**Fig 3d**).

Because tetanus immunity is long-lived and broadly prevalent in Singapore,^40^ we stimulated BTO with tetanus toxoid (TT) to assess recall humoral responses. ELISA data in **Fig 3e** showed that BTO elicited robust TT-specific IgG production, confirming that the system retained pre-existing memory of donors. Addition of a weak adjuvant, Alum, modestly boosted the titer of secreted TT-specific antibody. Overall, these results demonstrate that BTOs preserve cellular diversity and donor immune memory necessary for antigen-specific activation, GC formation, plasmablast differentiation, and antibody secretion. The BTO model can also discriminate different antigenic materials and offer a physiologically relevant *ex vivo* model to measure vaccine efficacy and strength of humoral recall immunity.

### BTOs mount robust responses against inactivated whole virus

To examine recall responses to complex antigenic repertoires, we challenged BTOs with inactivated influenza A (H1N1), a clinically relevant seasonal pathogens and vaccine target.^41,42^ Inactivated H1N1 stimulation triggered strong B cell activation and GC responses after 14 days post stimulation, including a significant increase in GC plasmablasts, indicating faithful recapitulation of B cell maturation to whole virus exposure in BTOs (**Fig 4a**, **Fig S4a**). Moreover, the inactivated H1N1 stimulation increased frequency of conventional dendritic cells (cDCs) and their maturation (**Fig S4b and c**), consistent with their role in bridging innate and adaptive immunity.^43^ The increase in cDCs suggested that the organoid microenvironment supported sensing of viral antigens, and mounting appropriate response through antigen-presenting cell recruitment and maturation. In line with these findings, H1N1-specific IgG levels rose modestly as shown by ELISA (**Fig 4b**). These results highlight the capacity of organoid models to support B cell differentiation and antibody production upon viral challenge.

**Figure 4.**
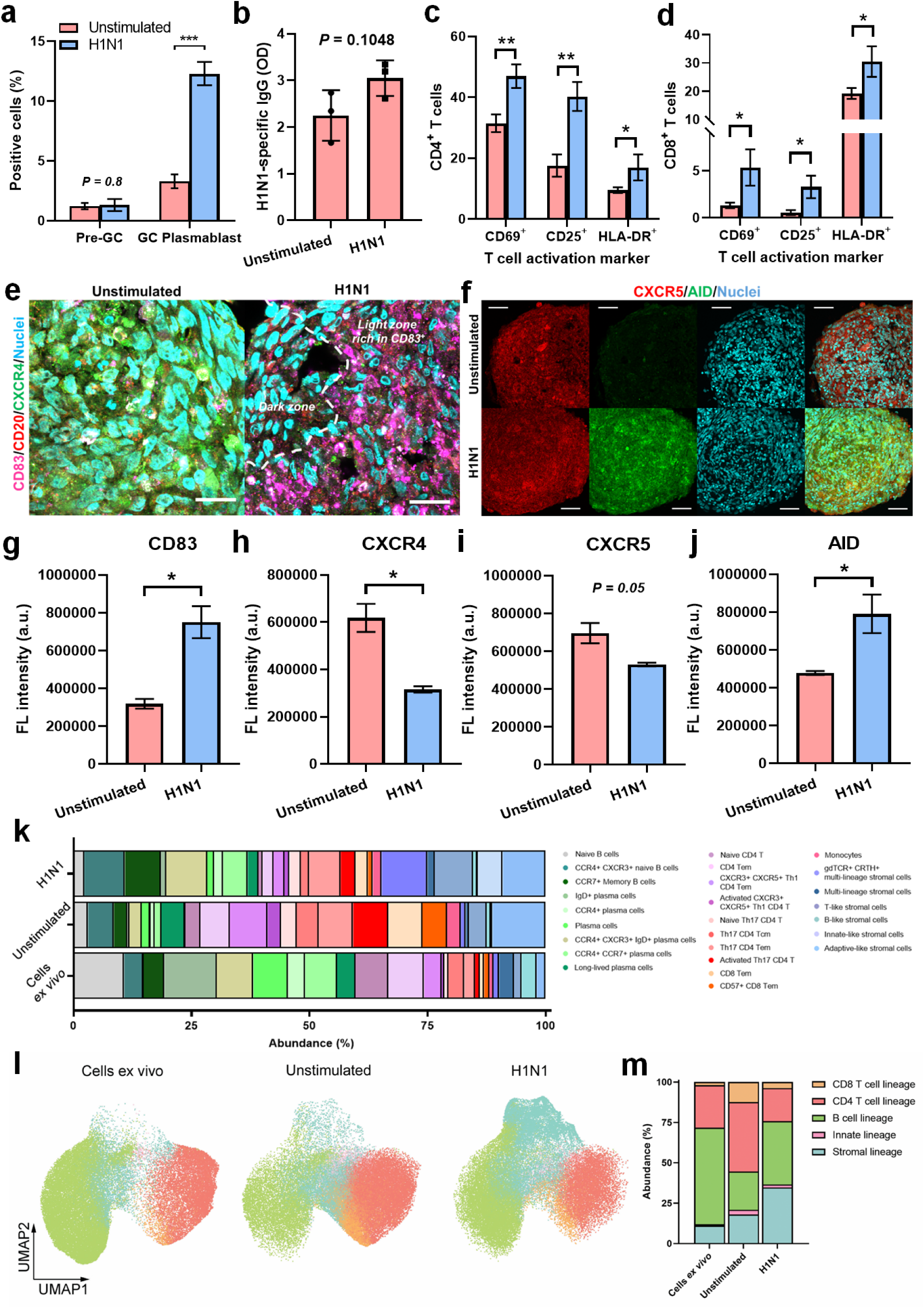
Tonsil organoids respond effectively to inactivated influenza A virus stimulation. **(a)** Bar graph of GC B cell responses after stimulation with inactivated H1N1 virus. **(b)** ELISA data of total antibody (IgG) production after stimulation. Bar graphs showing an increase of activated **(c)** CD4+ T cells and **(d)** CD8+ T cells after stimulation. Confocal microscopy images of tonsil organoid which represent **(e)** CD83 and CXCR4 expression corresponding to light- and dark zone in GC, respectively (scale bar = 20 µm), and **(f)** CXCR5 and AID expression corresponding to somatic hypermutation after stimulation (scale bar = 50 µm). Semi-quantification of **(g)** CD83 and **(h)** CXCR4 expression levels represent germinal center and **(i)** CXCR5 and **(j)** AID expression levels correspond to somatic hypermutation in BTO model after stimulation. **(k)** Bar plot showing multicellular dynamics of immune cells in tonsil organoid after stimulation (H1N1 organoid) compared to unstimulated (HD organoid) and cells *ex vivo*. **(l)** UMAP data and **(m)** bar graph showing clusters and relative abundance percentage of major immune cell subsets in H1N1 stimulated organoid compared to HD organoid and cells *ex vivo*. All data in **(k)** to **(m)** are based on cytometry by time of flight (CyTOF) analysis. All data are presented as mean ± S.D. Unpaired, two-tailed t-test was performed to calculate statistical significance. \**P* < 0.05; ** *P* < 0.01; \*\*\**P* < 0.001; no significance (n.s.), *P* > 0.05.

T cells in BTO were likewise responsive to viral stimulation. Flow cytometric data of T cell activation markers revealed significant increase in CD4⁺ T cell population (**Fig 4c**) to provide signals for promoting GC formation, affinity maturation, and plasma cell differentiation.^44^ Similarly, CD8⁺ T cells exhibited enhanced activation, indicating that the organoids are capable of eliciting cytotoxic T cell responses (**Fig 4d**). The coordinated activation of both CD4⁺ and CD8⁺ subsets demonstrated that BTO preserved important cellular crosstalk for adaptive immune defense against viral pathogens. Immunofluorescence images further provided spatial validation of these immune responses within the 3D organoid architecture. CD83 and CXCR4 expression pattern reflected light- and dark-zone organization of GC, respectively, served as a hallmark feature of active B cell maturation (**Fig 4e and Fig S5**). CXCR5 with activation-induced cytidine deaminase (AID) confirmed somatic hypermutation affinity maturation (**Fig 4f**). Semi-quantification showed significant upregulation of GC zonation markers and somatic hypermutation after H1N1 stimulation. Collectively, our data shows that the BTO model reproduces critical aspects of human adaptive immunity (**Fig 4g-j**).

Finally, CyTOF profiling compared the multicellular composition of H1N1-stimulated BTO with unstimulated BTO and freshly isolated tonsil cells. Stimulation shifted the cellular ecosystem toward activated B cell and T cell subsets (**Fig 4k**) and diversified stromal cell populations, suggesting stromal cells act as active participants rather than passive scaffolds. In lymphoid organs, FRCs and related stromal subsets organize network that guide lymphocyte and DC positioning to enable efficient antigen encounter and complex immune cell interactions.^45,46^ In **Fig 4l**, dimensionality reduction by UMAP demonstrated distinct clustering across *ex vivo* tonsil, unstimulated-, and H1N1-stimulated BTOs. The data highlights how viral challenge remodels immune cell compositions and architecture in BTO model, yet the diversity of immune cell populations was still preserved. The proportional cell distribution was slightly shifted, particularly with a relatively increased population of T cells and stromal subsets. Semi-quantification confirmed enrichment of adaptive lineages after H1N1 stimulation, possibly reflecting the transition from a basal state to an antigen-driven adaptive response (**Fig 4m**). Taken together, our data demonstrated that the BTO model reconstitute hallmarks of GC biology, including light/dark-zone architecture, somatic hypermutation, coordinated CD4⁺/CD8⁺ activation, while maintaining multicellular dynamics observed *in vivo*. These features position the BTO system as a physiologically relevant platform for investigating human antiviral immunity and for comparative evaluation of vaccine candidates in a controlled *in vitro* setting.

### Spatial analyses capture cell type-specific remodeling and cell-cell crosstalk after immune challenges

We used GeoMx Whole Transcriptome Atlas (WTA) spatial transcriptomic profiling to resolve cell type–specific responses to H1N1 in BTO. Regions of interest (ROIs) were segmented by lineage markers for B cells (CD20^+^), T cells (CD3^+^), and stromal cell subsets (PDPN^+^) with DNA counterstain as structural reference (**Fig 5a**). In unstimulated BTOs, CD20^+^ B cells were sparse to discrete clusters, whereas CD3^+^ T cells were distributed through the core of the organoid. In contrast, H1N1 stimulation produced more distinct B and T cell clusters, consistent with immune-stromal reorganization, GC-like compartmentalization, and enhanced crosstalk.

**Figure 5.**
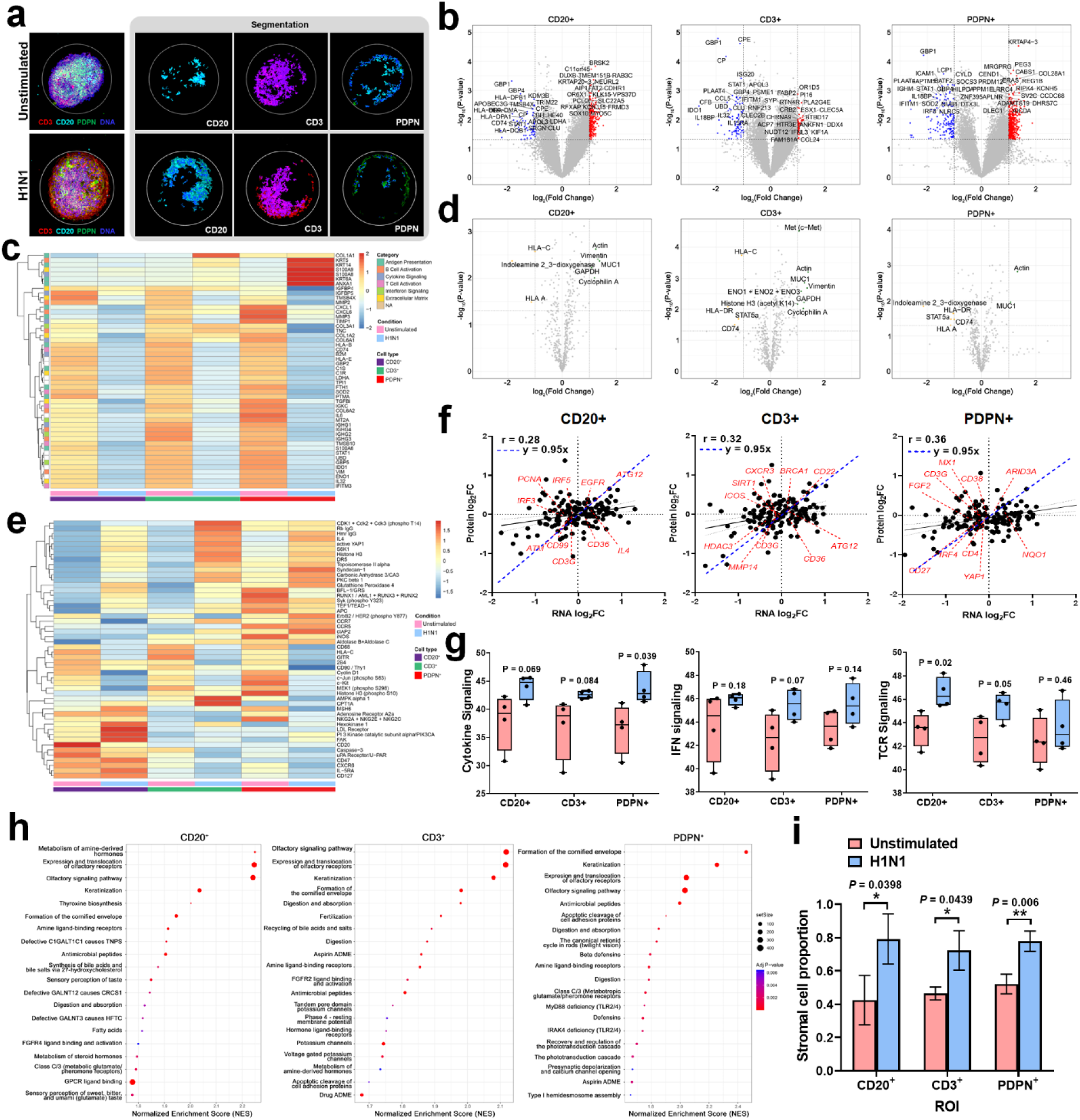
Digital spatial profiling revealed distinct transcriptomic and proteomic profiles of cell-type-specific responses in BTOs after immune challenge. **(a)** Representative images of regions of interest (ROIs) in unstimulated and H1N1 stimulated organoid. Fresh frozen organoid sections were stained with antibodies against CD20 (B cells), CD3 (T cells), and PDPN (stromal cells) to determine corresponding cells within ROIs. **(b)** Volcano plots showing differential gene expression analysis between unstimulated and H1N1 stimulated organoid in CD20+, CD3+, and PDPN+ compartment. **(c)** Unsupervised hierarchical clustering heatmap of top 50 differentially expressed RNA heatmap in BTO after stimulation. **(d)** Volcano plots representing protein expression profiles between unstimulated and H1N1 stimulated organoid in CD20+, CD3+, and PDPN+ compartment. **(e)** Unsupervised hierarchical clustering heatmap of protein targets in BTO after stimulation. **(f)** Scatter plots display RNA–protein correlation for selected targets within CD20⁺ B cell, CD3⁺ T cell, and PDPN⁺ stromal cell compartments between unstimulated and H1N1 stimulated organoids. **(g)** box plots showing major signaling pathways that have changed in organoid after stimulation, including cytokine-, IFN-, and TCR signaling. **(h)** Functional enrichment dot plots of Reactome pathways in CD20⁺, CD3⁺, and PDPN⁺ compartment, demonstrating distinct biological processes engaged by each cell population. **(i)** Transcriptional cell deconvolution data of stromal cell proportion in different segments (CD20⁺, CD3⁺, and PDPN⁺). Stromal cell populations are significantly increased in all segments after stimulation, indicating strong immune-to-stromal cell interaction during immunological responses. All data are presented as mean ± S.D. Multiple t-test was performed to calculate statistical significance. \**P* < 0.05; ** *P* < 0.01; \*\*\**P* < 0.001; no significance (n.s.), *P* > 0.05.

Differential expression analyses revealed broad transcriptional reprogramming across all cellular compartments in response to H1N1 stimulation. RNA volcano plots showed significantly upregulated (red dot) and downregulated (blue dot) genes in CD20⁺ B cells, CD3⁺ T cells, and PDPN⁺ stromal cells compared to unstimulated control (**Fig 5b**). In CD20⁺ B cell and CD3⁺ T cell regions, immune regulatory genes such as *GBP1, STAT1*, *HLA-DPB1*, *CCL5*, and *IDO1* were markedly downregulated, suggestive of viral evasion, while innate immune-related gene (*BRSK2*), metabolic and signaling-associated genes (e.g., *MYO5C* and *SOX10*), and inflammatory cytokine signaling gene (*RAB3C*) were markedly upregulated, indicating host antiviral responses within the BTO environment. In PDPN⁺ stromal ROIs, immuno-modulatory and chemokine-mediated transcripts like *ICAM1*, *GBP4, IRF8*, and *SOCS3* were broadly reduced, consistent with a regulatory adaptation to limit tissue inflammation, whereas genes linked to ECM remodeling (*COL28A1*, *ADAMTS19*, *PEG3*) and stromal differentiation (*AICDA*, *PEG3, REG1B*) were strongly upregulated, pointing to viral infection-driven matrix renewal and tissue repair in the stromal cell population.

The majority of transcripts in BTO whose expressions was significantly altered across CD20⁺ B cells, CD3⁺ T cells, and PDPN⁺ stromal regions after stimulation are presented in **Fig S6**. In **Fig 5c**, the RNA heatmap depicted hierarchical clustering of the top 50 differential expressed, immune-related genes that are up- or down-regulated across all cell types following viral exposure.

Expression of immune-response genes, including antigen presentation (e.g., *HLA-B*, *CD74*, *B2M*) and cytokine signaling mediators (*IL6*, *CXCL1*) was reduced. This broad suppression of innate and adaptive programs was consistent with influenza-mediated host shut-off mechanisms driven by viral NS1 protein, and/or a transition toward immune exhaustion or resolution. ^47–49^ In PDPN⁺ stromal cells after stimulation, structural genes (*COL1A1*, *KRT5*, *KRT6A*, and *KRT14*), inflammatory alarmins (*S100A8* and *S100A9*), and the anti-inflammatory mediator (*ANXA1*) were up-regulated. ECM dynamics in BTO was supported by significant shift in collagen gene profiles alongside with morphological changes (**Fig S7a**), with collagen-related transcripts up-regulated after H1N1 stimulation (**Fig S7b**), consistent with swelling of lymphoid tissue during viral infection. These data demonstrated stromal-driven tissue remodeling and stromal support of immune development in response H1N1. Spatial analyses thus revealed functional stromal patterns validated by alarmin release and tissue repair signaling amidst broad transcriptional suppression in immune compartments.

An immune-oncology proteome atlas (IPA) panel profiled over 570 proteins relevant to immune regulation, checkpoint signaling, and metabolic reprogramming, enabling proteomic readouts across immune and stromal compartments.^50^ Cytoskeletal and metabolic proteins (Actin, Vimentin, GAPDH, and Cyclophilin A), were significantly up-regulated in CD20^+^, CD3^+^ and PDPN^+^ ROIs, whereas immune-regulatory molecules (CD74, indoleamine 2,3-dioxygenase), antigen presentation-related molecules (HLA-A, HLA-C) and metabolic enzymes (ENO1/ENO2/ENO3) were significantly decreased after H1N1 stimulation (**Fig 5d**). These phenomena suggest adjustment in antigen-presenting capacity in immune-stromal compartments within BTO upon viral challenge, potentially reflecting immune resolution. The top 50 protein heatmap corroborated protein-level differences seen in volcano plots. H1N1 stimulation increased key inflammatory and signaling proteins (iNOS, cLAP2, CCR7, CCR5), particularly in CD3⁺ and PDPN⁺ regions (**Fig 5e**), indicating immune activation and tissue environmental responses. Concurrently, proteins involved in homeostasis, cell adhesion, and inflammatory metabolism (CD127, CXCR6, PIK3CA, and FAK) were up-regulated in CD20⁺ B cell compartments. Overall, these results highlight dynamic remodeling of immune and stromal proteomes during antiviral responses in the BTO model.

Notably, changes in protein amounts aligned well with transcriptomic trends from parallel RNA profiling. Scatter plots in **Fig 5f** illustrated gene-protein pair correlations, with most pairs showing concordant directionality and a subset diverging substantially due to potential post-transcriptional or post-translational regulation. For instance, upregulation of immune-related transcripts such as CD36 and IL4 in CD20^+^ B cell region and BRCA1, ATG12, and CD22 in CD3^+^ T cells exhibited strong directional correlation with their protein counterparts, implying to the immune activation, canonical signaling, and co-stimulatory pathways after viral stimulation. Meanwhile the YAP1/ARID3A and NQO1 correlated expression in PDPN^+^ stromal cells reflected stromal-immune interactions and interferon responses with minimum post-transcriptional modifications. These findings support the value of integrating transcriptomic and proteomic datasets to gather a more comprehensive understanding of immune-stromal compartments undergoing metabolic, structural, and antigen-presentation shifts in response to viral stimulation.

Gene set enrichment analysis (GSEA) provided insights into BTO responses to viral infection. Data in **Fig 5g** showed up-regulated pathways shared across all three different ROIs, including cytokine signaling, interferon responses, and T cell receptor pathways, possibly due to antiviral defense mechanisms, T cell activation and immune crosstalk with B cells. Reactome-based analyses detailed top enriched pathways in H1N1-stimulated BTO (**Fig 5h**). For example, CD20^+^ B cells showed up-regulated pathways associated with hormone metabolism, keratinization, and antimicrobial peptide production, related to metabolic adaptation, defense mechanism activation, and stress responses against viral infection. While enrichment in olfactory signaling pathways, fertilization-related processes, and ion channel regulation in CD3^+^ T cells demonstrated dynamic changes in membrane signaling, receptor activity, and cellular metabolism that may underpin T cell activation. In PDPN^+^ stromal cells, up-regulation of TLR-related signaling (including MyD88/IRAK4 axes) highlighted the engagement of stromal subsets with innate immune defense and tissue remodeling caused by infection. Transcriptional deconvolution further revealed increased proportion of stromal cells across all compartments after H1N1 stimulation (**Fig 5i**), supporting enhanced immune-stromal interactions and organoid structural remodeling to shape immune architecture and sustain adaptive responses.^21,51,52^ Taken together, the spatial multi-omic data supports that the BTO mounted a highly compartmentalized yet coordinated antiviral program, integrating distinct transcriptional and proteomic modules across immune and stromal niches.

### BTO models learn to recognize naïve antigens

Beyond retaining immune memory, it is important to demonstrate that the BTO model can be primed to recognize naïve antigens and recapitulate *de novo* immune responses.^9,50^ This function is essential for using the model to develop vaccine candidates and study novel pathogens or antigens to which human donors have not been previously exposed. We next assessed the BTO system for its ability to mount immune responses against naïve antigens, an ability not shown by published immune organoid models, except with R-phycoerythrin (R-PE) by Wagar et al.^12,13,16,17^ We stimulated the BTO model with R-PE, a foreign fluorescent protein to which all donors had no prior exposure. Flow cytometry detected a distinct population of R-PE-binding B cells, which was approximately 6.7-fold higher than in unstimulated controls (**Fig 6a-b**). This evidence hinted that BTO was capable of initiating antigen recognition even against wholly novel protein structures.

**Figure 6.**
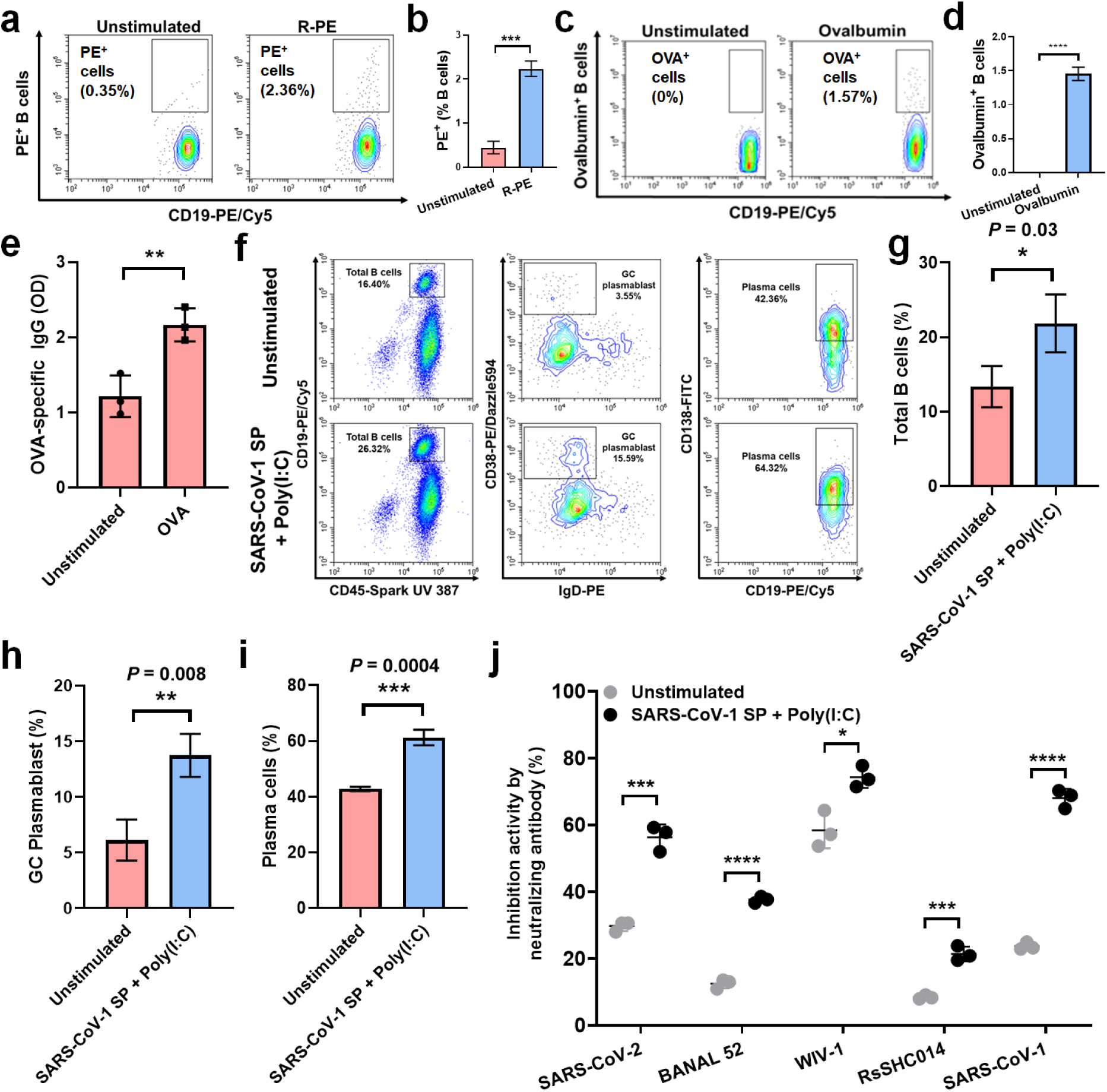
Tonsil organoids can learn to recognize naïve antigens. **(a)** Representative flow cytometry plots and **(b)** bar graph of B cell response towards R-phycoerythrin (R-PE) stimulation. **(c)** Representative flow cytometry plots and **(d)** bar graph of B cell response towards ovalbumin (Ova) stimulation. **(e)** ELISA data showing the capability of the BTO model to produce antigen-specific IgG production after naïve OVA stimulation. Representative flow cytometry staining **(f)** and bar graphs showing **(g)** total B cells, **(h)** GC B cell responses, and **(i)** plasma cell differentiation after SARS-CoV-1 spike protein stimulation in combination with poly(I:C) adjuvant. **(j)** Multiplex surrogate virus neutralization test (sVNT) analysis of tonsil organoid supernatant against 6 different sarbecoviruses. Naïve stimulation with SARS-CoV-1 spike protein showed the ability of BTO to produce cross-clade neutralizing antibody against SARS-CoV-1 virus family. All data are presented as mean ± S.D. Unpaired, two-tailed t-test was performed to calculate statistical significance. \**P* < 0.05; ** *P* < 0.01; \*\*\**P* < 0.001; no significance (n.s.), *P* > 0.05.

We then used a gold-standard naïve antigen, ovalbumin (OVA), widely used to study antigen presentation, T cell activation, and humoral immune responses due to its well-characterized structure, epitopes, and processing pathways,^53^ **Fig 6c-d** showed that OVA-specific B cells were readily detected after OVA stimulation, while the signal was absent in unstimulated conditions. Importantly, functional readouts of this antigen recognition were reflected by the production of OVA-specific IgG antibody measured by ELISA in stimulated BTO supernatant (**Fig 6e**). These findings showed that the BTO model supported a complete humoral response, from naïve antigen uptake and processing to secretion of antigen-specific immunoglobulins.

To further demonstrate translational potential of BTO platform, organoids were challenged with a naïve virus. Severe acute respiratory syndrome coronavirus 1 (SARS-CoV-1) is a zoonotic betacoronavirus responsible for the 2002–2003 outbreak, spreading to more than 30 countries, including Singapore, with a ∼10% case fatality rate. ^54^ Based on age and serology, our donors were immunologically naïve to SARS-CoV-1, as the virus had not circulated in humans since the outbreak was contained in 2004. We previously reported that human subjects infected with SARS-CoV-1 elicited broadly neutralizing antibodies against cross-clade viruses, a feature not replicated in 2D immune cell culture.^55^ The BTOs were stimulated with SARS-CoV-1 spike protein as naïve viral subunit antigen together with the TLR3 agonist poly(I:C), as an immune adjuvant. Flow cytometry staining and quantification revealed robust expansion of total B cells, significant induction of GC plasmablasts, and downstream differentiation into plasma cells (**Fig 6f–i**). This increase in B cell subsets recapitulated the stepwise *in vivo* progression of activation, affinity maturation, and effector differentiation, suggesting that the BTO microenvironment provided cues necessary for antigen-driven B cell maturation against naïve infection.

We next assessed functional relevance using a multiplex surrogate virus neutralization test (sVNT) to detect antibodies in the supernatants using previously described protocol.^56^ Stimulation with SARS-CoV-1 spike protein elicited antibodies with cross-clade neutralizing capacity that inhibited not only the cognate antigen but also spike proteins from other sarbecoviruses (**Fig 6j**). These data underscored the ability of BTO to mount *de novo* humoral responses against naïve viral antigens and recapitulates a phenomenon observed uniquely in SARS-CoV-1 infected humans, i.e. generation of a broad spectrum of neutralizing antibodies targeting conserved epitopes across related viral species. Our results support the use of BTO as a pre-clinical platform to evaluate primary adaptive responses to novel antigens in vaccine development and adjuvant testing.

## Discussion

In this study, we developed a BTO model using an original strategy that combined autologous stromal cells with immune cells to recapitulate core features of a secondary lymphoid tissue. Stromal cells are essential sentinels in immune organs as they provide scaffold, direct immune cell trafficking, deliver survival and differentiation cues, regulate immune responses, and sustain surveillance.^57,58^ Yet, prior immune organoid models did not define or characterize stromal roles. Here, BTO maintained structural integrity, multicellular diversity, and functional GC biology by restoring a FRC–like stromal niche that supported antigen-specific B and T cell responses. Optimizing the proportion of PDPN⁺ stromal cells stabilize organoid morphology and facilitate immune-stromal crosstalk, whereas insufficient stromal support led to architectural collapsed or loose aggregates,^12^ underscoring the requirement for cytoskeletal and matrix programs to organize immune interactions. Our CyTOF data showed that the BTO preserved major lineages with subset diversity comparable to freshly isolated *ex vivo* tissue-derived cells. Compared to the transwell organoid model, the BTO model sustained a richer stromal heterogeneity and a more balanced repertoire of immune cells, underscoring stromal engineering as a prerequisite for physiological fidelity.

Functionally, BTO mounted coordinated responses to canonical immune stimuli, i.e., PHA and LPS to sustain survival, and R848/IL-2 cocktail to trigger B and T cell activation. All of these demonstrated integration of innate cues with adaptive activation in the BTO model. Moreover, the organoid deployed effective recall antigen-specific immunity. Stimulating BTOs with COVID-19- associated viral subunits revealed a clinically concordant hierarchy of immunogenicity (RBD < live virus < spike < mRNA vaccine) for plasmablast generation, plasma cell differentiation, and IgG production, which never been studied in other reported immune organoid models. Whole-virus H1N1 challenge of BTO further capture complex immune dynamics that is comparable to previous studies. ^12,13,17^ Viral exposure induced GC responses, H1N1-specific IgG production, and activation of CD4⁺ helper and CD8⁺ cytotoxic T cells. Confocal imaging revealed parallel expressions of GC light- and dark-zone marker (CD83/CXCR4) and hallmarks of somatic hypermutation and affinity maturation (AID/CXCR5) in stimulated BTO. Additionally, multicellular profiling showed enrichment of adaptive immune subsets with diversification of stromal cells, suggesting that stromal populations actively participated in antigen capture, chemokine guidance, and tissue remodeling - roles ascribed to FRC-like networks *in vivo*.

Spatial transcriptomic and proteomic analyses resolved compartment-specific remodeling within CD20⁺ B cells, CD3⁺ T cells, and PDPN⁺ stromal cells. Viral challenge suppressed antigen presentation and cytokine programs in immune regions, compatible with peak activation transitioning into an early resolution or viral host-shutoff phase. Meanwhile, the stromal cell compartment upregulated ECM-related genes and alarmins, which indicated dual roles in tissue remodeling and inflammatory regulation to support trafficking, mechanical integrity, and controlled resolution of inflammation in stimulated BTO. Proteomic profiling partially mirrored RNA trends and revealed complementary shifts in cytoskeletal, metabolic, and signaling proteins (e.g., iNOS, CCR5, CCR7), consistent with post-transcriptional regulation in BTO. Taken together, the results reinforce that stromal cells are not passive scaffolds, but active regulators of immune activation and resolution in organoid confinement.

Most importantly, BTO demonstrated *de novo* responses to naïve antigens beyond recall immunity. SARS-CoV-1 spike protein plus poly(I:C) adjuvant induced GC formation, plasma cell differentiation, and production of cross-clade neutralizing antibody against multiple sarbecoviruses. This demonstrates the potential of our BTO model to reveal conserved epitope targeting in SARS-CoV-1 infected humans and supports its value for evaluating novel antigens aimed for pan-variant and pan-lineage protection in pandemic preparedness. With the passage of the FDA Modernization Act 2.0, reliance on animal models is no longer mandated for preclinical drug testing, opening opportunities for the timeliness and translational potential of our BTO model. In summary, we believe that the BTO model is uniquely poised to make a major contribution for *in vitro* modeling of human immune responses including applications in vaccine development, testing of material immunogenicity and immune mechanisms that are otherwise hard to study using PBMCs.

Nonetheless, we acknowledge several limitations of our work that warrants more development. Specialized stromal subsets (e.g., follicular dendritic cells) and epithelial cells remain under-characterized and should be further incorporated into the model to enhance physiological authenticity. The absence of vascular or lymphatic flow may constrain antigen transport and effector cell egression in BTO system. Addition of dynamic perfusion could extend culture longevity and mechanistic reach for human immunology and preclinical immunotherapeutics testing.

## Materials & Methods

### Informed consent and sample collection

Human tonsils from consented patients undergoing tonsillectomy were collected in accordance with the National University of Singapore Institutional Review Board (NUS-IRB-2022-475). The participants were adults with age range from 21 – 39 years who went through surgery for obstructive sleep apnea or hypertrophy, and all tissues were typically healthy. Whole tonsils were collected immediately after surgery and then immersed in Ham’s F12 medium (Gibco) supplemented with 1× Normocin (InvivoGen) and 1% Penicillin-Streptomycin for 1 h at 4 °C for decontaminating steps. Tissues were subsequently rinsed with PBS prior to the tissue processing.

### Tissue processing

Tonsil tissue was diced into small pieces, followed with enzymatic digestion with fresh RPMI medium consisting of Collagenase type II (Invitrogen), Dispase (Sigma), and DNAseI (Sigma) with concentration at 200 U/ml, 2.4 mg/ml, and 0.3 mg/ml respectively. Minced tissue was enzymatically dissociated within 2 h at 37 °C in shaking water bath. Right after, cell suspension will be passed through a 100 µm and 70 µm cell strainer subsequently, and tissue debris was reduced by debris removal solution (Miltenyi) according to manufacturer’s protocol. Red blood cells were reduced by 1× RBC lysis buffer (Invitrogen). Cells were then washed with complete RPMI medium supplemented with Glutamax, 10% FBS, 1× non-essential amino acids, 1× sodium pyruvate, 1× insulin/transferrin/selenium (Gibco), 1× Normocin (InvivoGen), and 1% Penicillin-Streptomycin. Cell counting was performed, followed by cryopreservation with Fetal Bovine Serum (FBS) and 10% DMSO. Tonsil cells were kept at −80 °C freezer (short-term) or −196 °C liquid nitrogen (long-term) until use.

### Tonsil stromal cells (TSCs) isolation

Thawed tonsil cell stocks were plated on tissue culture flask and floating lymphocytes were removed after 72 hr, and the remaining adherent cells were cultured for 7-10 days. After the cells reached 80-90% confluency, they were detached with 0.25% Trypsin-EDTA for 5 min for cell passaging. The cells were characterized with flow cytometry and immunofluorescence analysis to check for positive expression of mesenchymal stem cell markers (CD44, CD73, CD90, and CD105), fibroblastic reticular cell (FRC) marker (Podoplanin/PDPN), and negative expression of blood cell markers (CD34), endothelial cell marker (CD31), and epithelial cell markers (CD324 or E-cadherin). TSCs were cultured in DMEM F/12 medium (Gibco) supplemented with 10% FBS and 1% Penicillin-Streptomycin. Cells were enumerated and divided into 1×10^6^ cells in each vial prior the cryopreservation.

### Organoid culture

Tonsil organoids were cultured using hanging drop (HD) method by making cell cocktails into droplets consisting of tonsil cell mixtures (TCMs) and TSCs at the optimum ratio. Thawed TCMs and TSCs were mixed in a 15 ml Falcon tube and centrifuged at 400 × g for 5 min. Cell pellet was resuspended in desired volume of complete RPMI medium following on the TCM:TSC ratio. A 10 µl of cell cocktail were made into a droplet on tissue culture dish lid and cultured under standard culture conditions (37°C, 5% CO_2_). Spherical aggregates observed after 48h were transferred into a multi-well tissue culture plate prior the antigen stimulation. Organoids were stimulated with various antigens as shown in **Table S2**. In terms of live virus infection, the organoids were were infected with 0.1 multiplicity of infection (MOI) of SARS-CoV-2 Omicron BA.2.86 subvariant. The infected organoids were incubated for 7 days at standard culture conditions. Prior the flow cytometry analysis, the dissociated organoids were fixed with 10% Neutral Buffered Formalin solution.

### Flow cytometry analysis

Organoids were dissociated into single cells with 0.25% Trypsin-EDTA. Cells were then washed with FACS buffer (PBS + 0.1% BSA + 2 mM EDTA). Cells were stained at 4 °C with anti-human antibodies in the presence of live/dead stain (Invitrogen). All antibodies were purchased from BioLegend as follows: CD45-Spark UV 387, CD19-PE/Cy5, CD27-PE/Cy7, CD27-BV711, CD38-PE/Dazzle 594, CD138-FITC, IgD-PE, IgM-Pacific blue, IgG-APC/Fire 750, CD3-APC, CD3-Alexa Fluor 700, CD4-PE/Cy7, CD8-BV605, CD25-BV750, CD69-BV421, HLA-DR-FITC, CD14-PE/Cy7, CD11c-PE, CD80-Alexa Fluor 647, CD83-BV421, CD40-FITC, CD123-BV711, CD123-Pacific blue, IL-2-BV421, and IFN-γ-BV711. Flow cytometry analysis was performed with CytoFLEX instrument (Beckman Coulter). For stromal fibroblastic cells characterization, the flow cytometric antibodies were as follows: BD Biosciences: CD73-PE, CD90-PE, CD34-PE, CD45-PE, CD14-PE, E-cadherin-Alexa Fluor 488, Biolegend: CD31, Podoplanin, Thermo Fisher: CD105-PE.

### Immunoglobulin detection by ELISA

Organoid culture supernatants were collected for ELISA assay. Total IgG detection was performed by ELISA kit (Thermo Scientific) following the manufacturer’s protocols. For tetanus toxoid-specific IgG detection, culture supernatants were assayed by human tetanus toxoid IgG ELISA kit (Creative Diagnostics) following the manufacturer’s instructions.

### Immunofluorescence analysis

Organoid samples were fixed with 10% neutral-buffered formalin (4 °C, overnight) and washed three times with PBS. Samples were then immersed in 30% sucrose solution (4 °C, overnight) and then embedded into 4% low-melting point agarose gel. Agarose-embedded organoids were added into a warm OCT compound in a cryomold and snap frozen on dry ice. Embedded samples were sectioned at 25 µm and collected onto poly-L-lysine coated glass slides. Sections were washed with PBS-T (PBS + 0.4% Triton X-100) and rehydrated with blocking buffer (PBS-T + 5% BSA) for 30 min. Sections were stained with primary unconjugated antibodies for 3 h at room temperature, followed with secondary antibody staining for 1 h at room temperature. Stained sections were further stained with primary direct conjugated antibodies for 3 h at room temperature (only if applicable). All primary antibodies were listed as follows: Biolegend: CD3-Alexa Fluor 488 (UCHT1; 1:100), CD20-Alexa Fluor 594 (2H7; 1:100), CXCR4 (12G5; 1:100), Podoplanin (NC-08; 1:250), CD3 (OKT3; 1:100), CD19 (HIB19; 1:100), CD45 (HI30; 1:100), Sigma-Aldrich: CD83 (Polyclonal; 1:200), CXCR5 (Polyclonal; 1:100), Thermo Fisher: CD20-eFluor 660 (L26; 1:100), AID (ZA001; 1:200), Abcam: Phalloidin-iFluor 647 (1:1000), Merck Millipore: Vimentin (V9, 1:100). All secondary antibodies were purchased from Thermo Fisher with dilutions as follows: Goat anti-Rabbit IgG (H+L) Cross-Adsorbed Secondary Antibody, Alexa Fluor 488 (1:200), Goat anti-Rabbit IgG (H+L) Highly Cross-Adsorbed Secondary Antibody, Alexa Fluor 594 (1:200), Goat anti-Mouse IgG (H+L) Highly Cross-Adsorbed Secondary Antibody, Alexa Fluor Plus 488 (1:200), Goat anti-Mouse IgG (H+L) Highly Cross-Adsorbed Secondary Antibody, Alexa Fluor Plus 594 (1:200), Goat anti-Rat IgG (H+L) Highly Cross-Adsorbed Secondary Antibody, Alexa Fluor Plus 488 (1:200), and Goat anti-Rat IgG (H+L) Highly Cross-Adsorbed Secondary Antibody, Alexa Fluor Plus 594 (1:200). Stained slides were counterstained with Hoechst 33342 (1 µg/ml) to visualize the nucleus. Imaging was performed using Nikon ECLIPSE Ti2 fluorescence inverted microscope and Olympus FV3000 Confocal Microscope. Images were processed with ImageJ software.

### Surrogate virus neutralization tests (sVNTs)

For multiplex sVNTs, we adapted the sVNT using the Luminex platform. AviTag-biotinylated receptor-binding domain (RBD) proteins from 10 different sarbecoviruses were coated on MagPlex-Avidin microspheres (Luminex) at 5 μg per 1 million beads. RBD-coated microspheres (600 beads per antigen) were preincubated with culture supernatant at a final dilution of 1:20 or greater for 1 hour at 37°C with 800 rpm agitation. After 1 hour of incubation, 50 μl of phycoerythrin (PE)–conjugated human angiotensin-converting enzyme 2 (ACE2) (hACE2; 1 μg per milliliter; GenScript) was added to the well and incubated for 1 hour at 37°C with agitation, followed by two washes with 1% bovine serum albumin in phosphate-buffered saline (PBS). The final readings were acquired with the use of the MAGPIX system (Luminex).

### Cytometry by time of flight (CyTOF)

The 31-plex antibody cocktail was prepared on the day of cell staining (see **Table S3**). Frozen cells were thawed and plated at 3 × 10^6^ in a 96-well U bottom plate, washed in cold PBS buffer and incubated with 200 μm Cisplatin on ice for 5 minutes. Cells were washed in CyFACS buffer then pre-stained with anti-CD27, anti-161, anti-CCR7, and anti-CD14 in 50 μL reaction volume for 30mins at 30°C. After pre-staining, cells were washed in cold CyFACS buffer and stained with fluorochrome conjugated anti-TCR-γ/δ for 30mins at room temperature. Then, the cells were washed with CyFACS buffer and resuspended with metal isotope-labeled surface antibodies at room temperature. After 30 minutes, cells were washed with CyFACS buffer, followed by PBS and fixed with 2 % v/v paraformaldehyde (PFA) at 4°C overnight. On the next day, PFA was removed by washing in CyFACS buffer and Cell-ID Intercalator-Ir (Standard Biotools) in in 2% PFA/PBS for 20 mins at room temperature. After 20 minutes, cells were washed twice with cold CyFACS. Immediately before mass cytometry acquisition, cells were washed twice with MilliQ water, filtered and diluted to a final concentration of 0.6 × 10^6^ cells/mL. Prior to mass cytometry acquisition, 10 % v/v of EQ Four Element Calibration Beads (Standard Biotools) was added, and samples were acquired on a Helios Mass Cytometer at an event rate of< 400 events per second.

### GeoMx Whole Transcriptomics Atlas (WTA) + IO Proteome Atlas (IPA)

Fresh frozen organoid samples were sectioned at 6 μm thickness and stored at −80 °C until use. Prior to beginning the workflow, slides were processed according to Appendix II of NanoString manual MAN-10150-06. We then followed the NanoString guidelines outlined in MAN-10158-05, incorporating steps from MAN-1050 through the Proteinase K treatment, with the concentration adjusted to 0.1 μg/mL. From the overnight hybridization step onward, the protocol reverted to MAN-10158-05. During primary antibody incubation, samples were stained for CD20 (Abcam, ab198943), CD3 (Novus Bio, NBP2-54392AF594), and PDPN (Abcam, ab196515) antibodies to identify B cells, T cells, and stromal cells subsequently. Samples were prepared according to Nanostring’s procedure and then loaded into the DSP instrument for selection of regions of interest. In situ hybridization of RNA-directed DNA oligo probes (NanoString Human Whole Transcriptome Atlas, 18,677 genes) and simultaneous protein detection using antibody oligo conjugates from the Immuno-Protein Atlas (IPA, 570 proteins) were performed on the same slide according to the manufacturer’s protocol. GeoMx data were analyzed using Rstudio with R package (v.4.4.3). Quality check and preprocessing of the data were done using the standard parameters as recommended by Nanostring (https://github.com/Nanostring-Biostats/GeomxTools). Quartile 3 normalization was used to normalize data. Differential gene expression analysis between H1N1-stimulated and unstimulated organoid was conducted with Student’s T-test in R. Gene set enrichment analysis (GSEA) was performed using ClusterProfiler in R package (v.4.16.0) based on Reactome database. Data were plotted using ggplot2 R package (v.3.5.2).

### Statistical analysis

All data were presented as mean ± standard deviation (S.D.) with three replications. The statistical significance between the two groups was determined using the unpaired Student’s t-test by GraphPad Prism 8 Software. Statistical significances in more than two groups will be analyzed using a one-way analysis of variance (ANOVA) with Tukey’s post hoc test by GraphPad Prism 8 Software. A significant difference was marked as * (p < 0.05), **(p < 0.01), ***(p < 0.001), and ****(p < 0.0001).

## Supporting Information

Supporting Information is available from …

## Acknowledgements

This research was supported by the NUS Presidential Young Professorship, Ministry of Education (MOE) Tier 1 (Project #: 23-0968-P0001), National Medical Research Council Open Fund Investigator Research Grant (Application ID: OFIRG24jul-0076), iHT OOE award, Joint NCIS Centre Grant and NUS Centre for Cancer Research (N2CR) Seed Funding Programme, TREX Grant, PCM Seed Grant, and PREPARE Strategic Open Grant Call (Vaccines & Therapeutics Co-Operative Programme) (Project #: PREPARE-CS1-2024-014). This project was also supported by the WTA+IPA grant jointly awarded by NanoString Technologies and Next Level Genomics (NLG), NUHS Seed Funding (NUHSRO/2022/060/RO5+6/Seed-Mar/05) and NUSMed BSL3 S3RT Grant. Spatial multiomic profiling was performed on the GeoMx Digital Spatial Profiler (DSP) platform using the Whole Transcriptome Atlas (WTA) and Immuno-Protein Atlas (IPA) assays. We thank Dr. Ufuk Degirmenci for his close collaboration and support in this study. This work was awarded second place in the NanoString/NLG WTA+IPA grant competition. We also thank the NUS Infectious Diseases Translational Research Programme (IDTRP) and NUS Medicine BSL-3 Core Facility team for their funding and operational support in obtaining the SARS-CoV-2 for live virus experiments.

## Authors Contributions

I.R.S., D.E.K.C., Y.C.F.S., G.J.D.S, and A.T. conceived the project and experiments. I.R.S., Z.Z.R.L., K.S.T., J.J.H.C., G.G.K.N., and C.W.T. performed experiments. I.R.S., H.S.C., and N.S.T. analyzed data. I.R.S. and A.T. wrote the manuscript. A.T. supervised the project. All authors discussed the results and manuscript. Please refer to **Table S4** for a comprehensive breakdown of the authors’ contributions.

