## Supplementary Information for "Bioengineered human tonsil organoids as an immuno-engineering platform for evaluating immune functions"

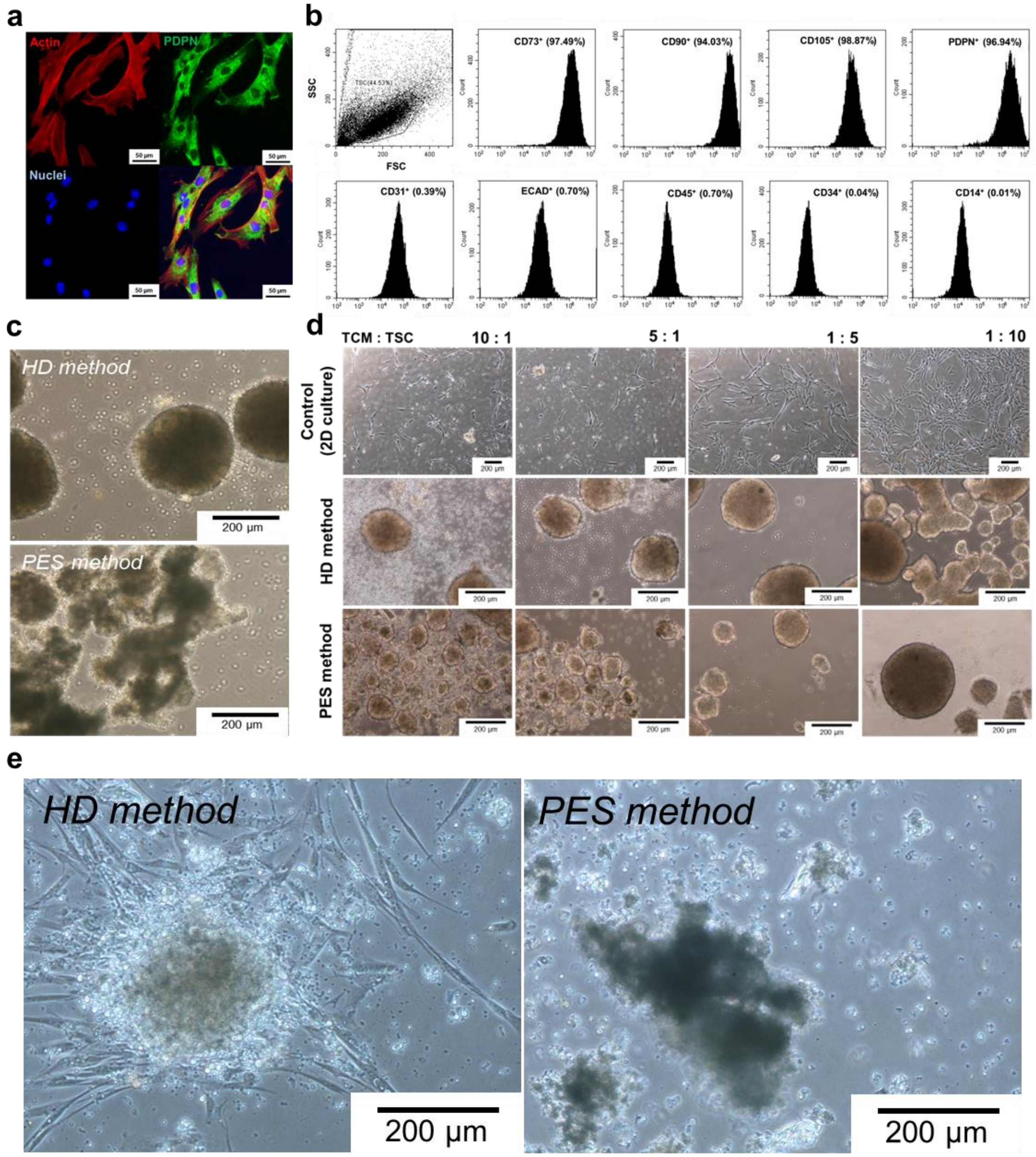

**Figure S1.** Autologous tonsil organoid culture characterizations. **(a)** Immunofluorescence images and **(b)** flow cytometry data of tonsil stromal cell characterizations showing high purity of cell. **(c)** Microscopic images comparing HD- and PES-based tonsil organoids. **(d)** Optimization of tonsil cell mixture and tonsil stromal cell (TCM:TSC) ratio for tonsil organoid showing that a minimum of 10:1 ratio was needed to generate organoids with controllable morphologies. **(e)** Microscopic image of organoid culture in 50:1 ratio with HD method (left) and PES method (right).

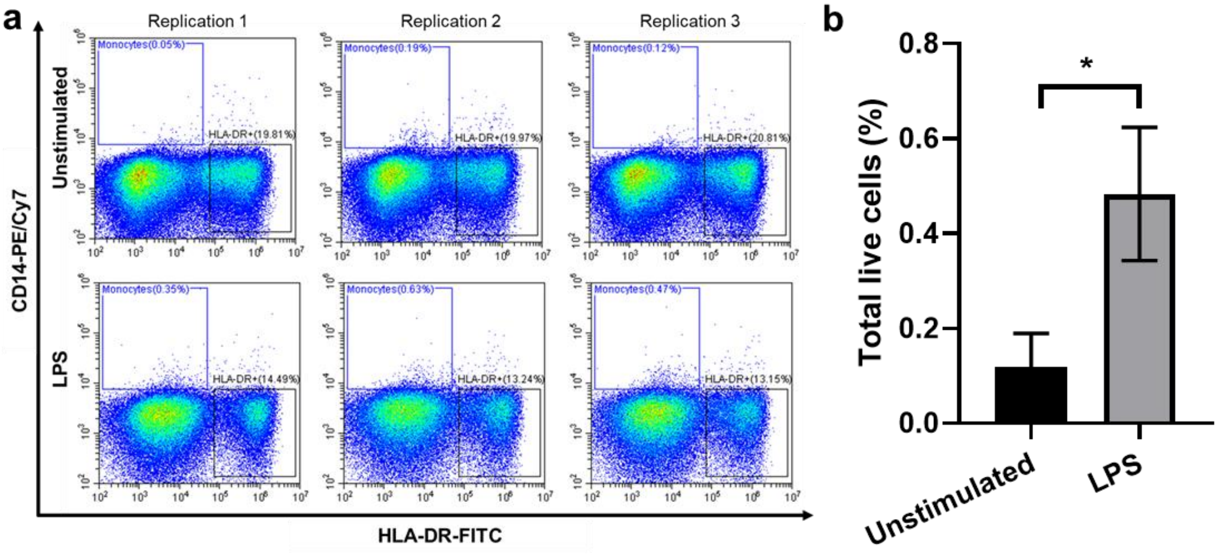

**Figure S2.** Monocytes population in BTOs after lipopolysaccharide (LPS) stimulation at concentration of 50 mg/ml. All data are presented as mean ± S.D. Unpaired, two-tailed t-test was performed to calculate statistical significance. *$P < 0.05$.

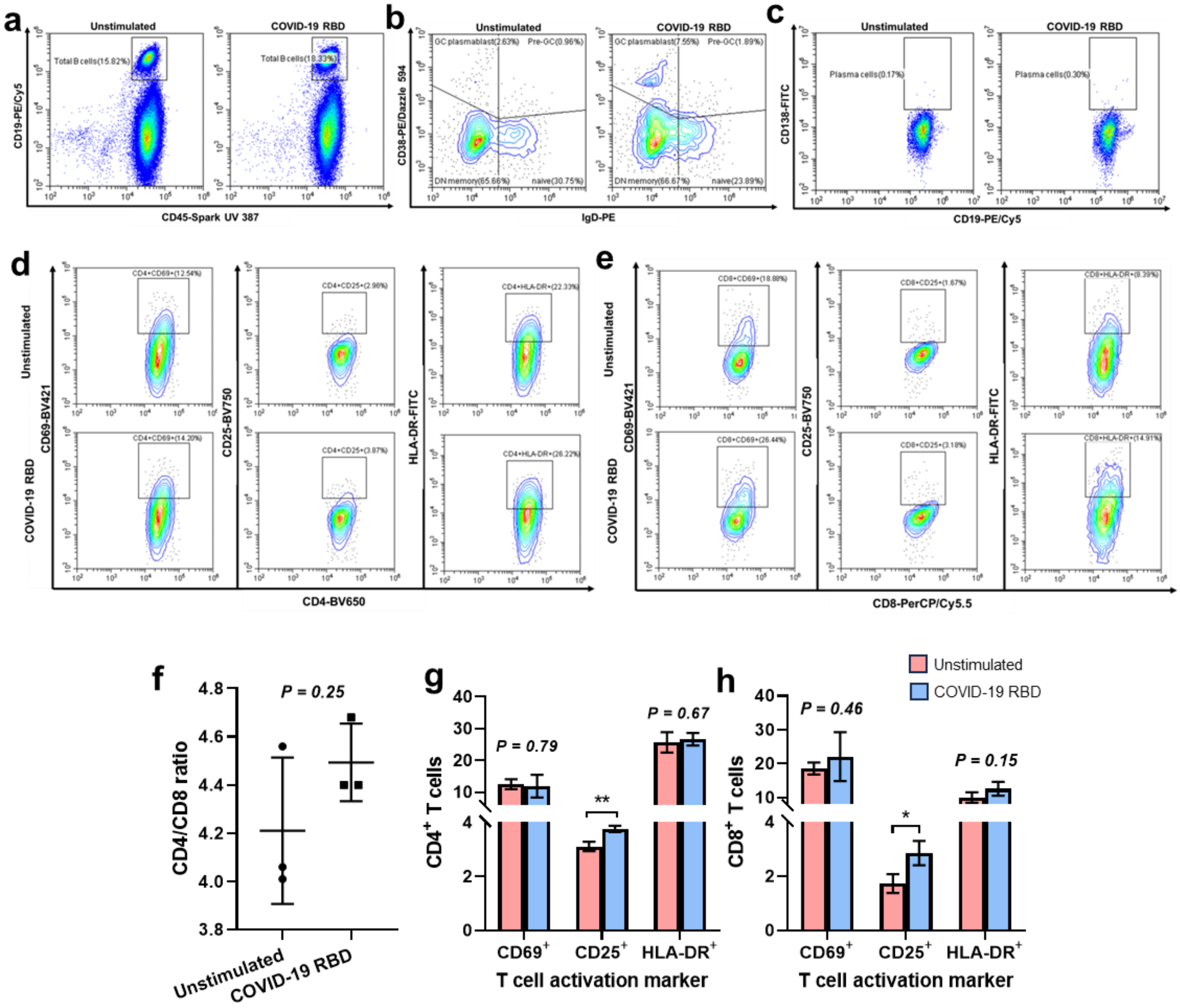

**Figure S3.** BTO model mounted adaptive immune responses after stimulation with COVID-19 receptor binding domain (RBD) protein. Representative flow cytometry plots of **(a)** Total B cells, **(b)** GC B cell responses, and **(c)** plasma cell differentiation after stimulation, indicating a recall humoral immunity in stimulated organoid. Tonsil organoid stimulation also led to cellular immune responses as shown by T cell aspects. Representative flow cytometry plots of T cell activation markers in **(d)** CD4 T cells and **(e)** CD8 T cells after stimulation. **(f)** Bar graph showing CD4 and CD8 T cell ratio in stimulated organoids. Bar graphs showing T cell activation marker expression of **(g)** CD4 T cells and **(h)** CD8 T cells after stimulation. All data are presented as mean ± S.D. Unpaired, two-tailed t-test was performed to calculate statistical significance. *$P < 0.05$; ** $P < 0.01$; no significance (n.s.), $P > 0.05$.

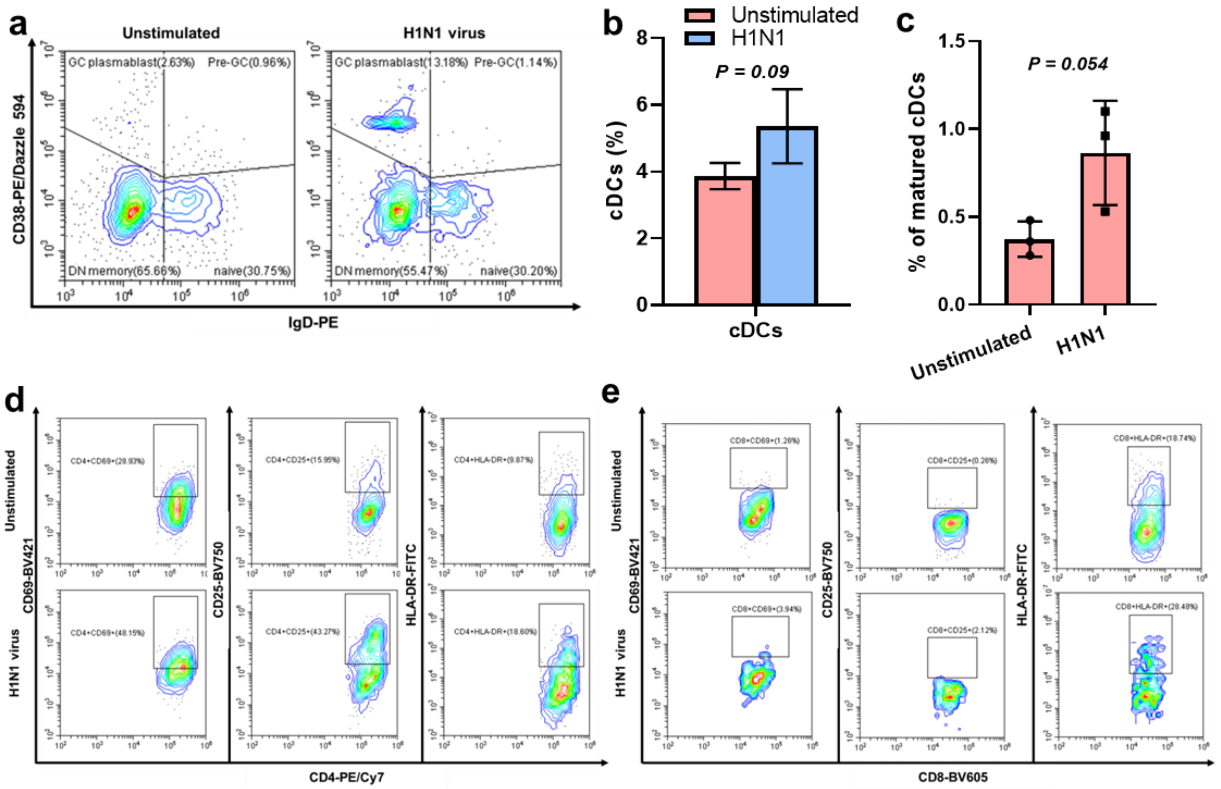

**Figure S4.** **(a)** Representative flow cytometry staining of GC B cell responses after inactivated H1N1 virus stimulation. **(b)** Percentage of cDCs and **(c)** their maturation level in tonsil organoid throughout the immune responses. Representative flow cytometry staining of T cell activation markers in **(d)** $CD4^+$ T cells and **(e)** $CD8^+$ T cells after stimulation. All data are presented as mean ± S.D. Unpaired, two-tailed t-test was performed to calculate statistical significance. *$P < 0.05$; no significance (n.s.), $P > 0.05$.

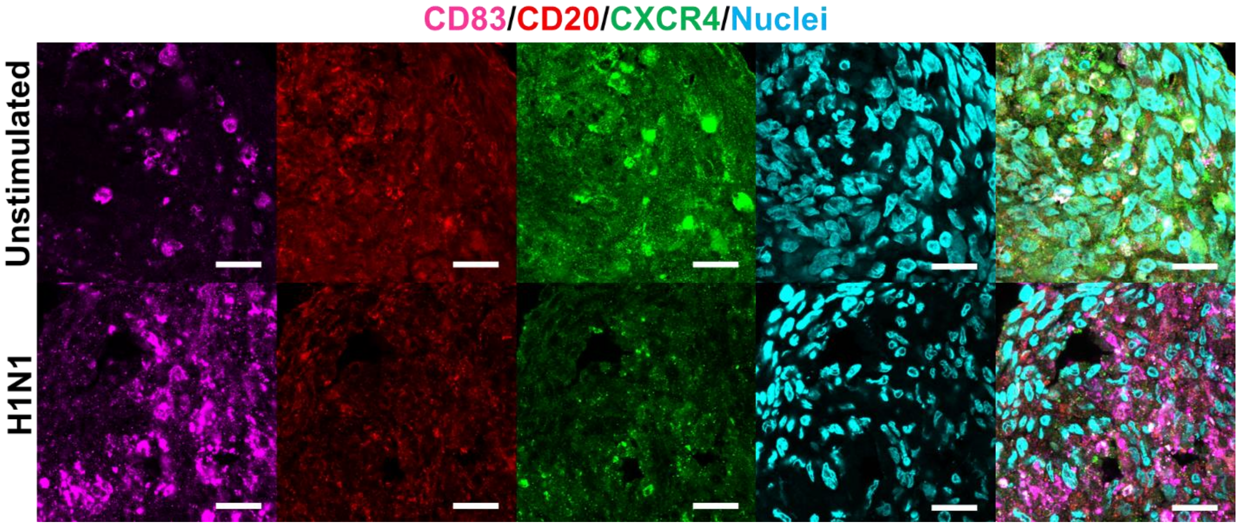

Figure S5. Confocal microscopy images of tonsil organoid which represent CD20, CD83, and CXCR4 expression corresponding to light- and dark zone in GC, respectively. Scale bar = 20 µm.

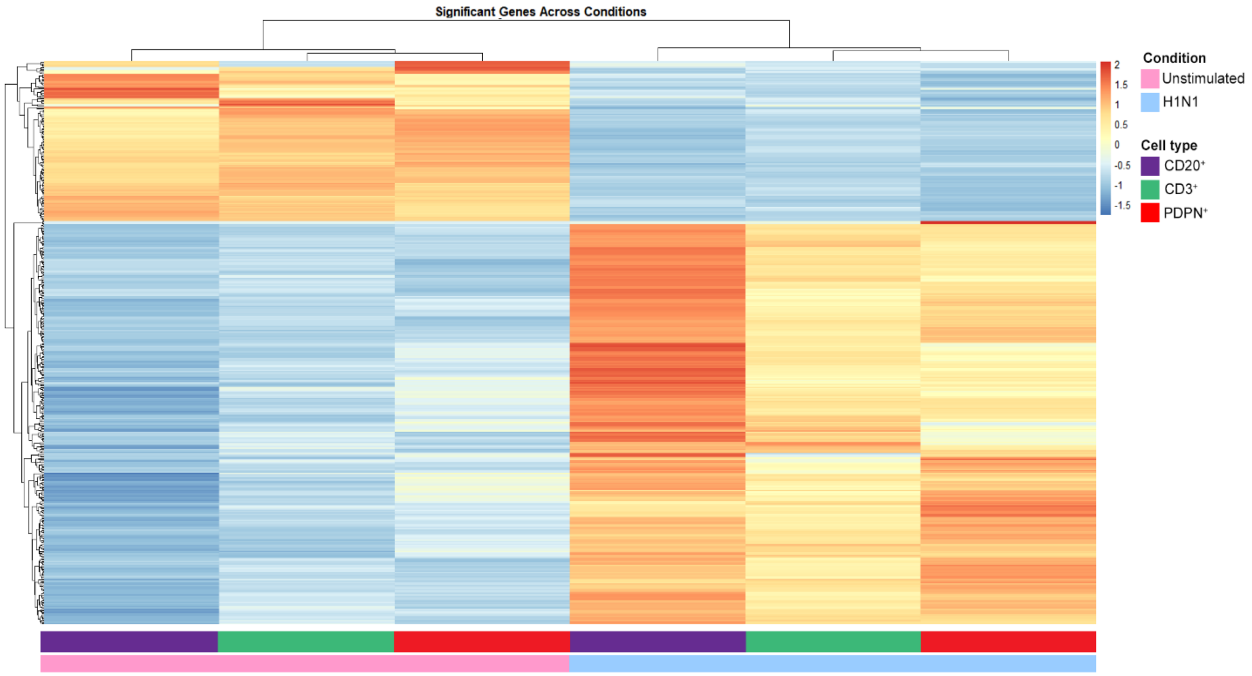

**Figure S6.** Unsupervised hierarchical clustering heatmap of all significant differentially expressed RNA heatmap in BTO after stimulation.

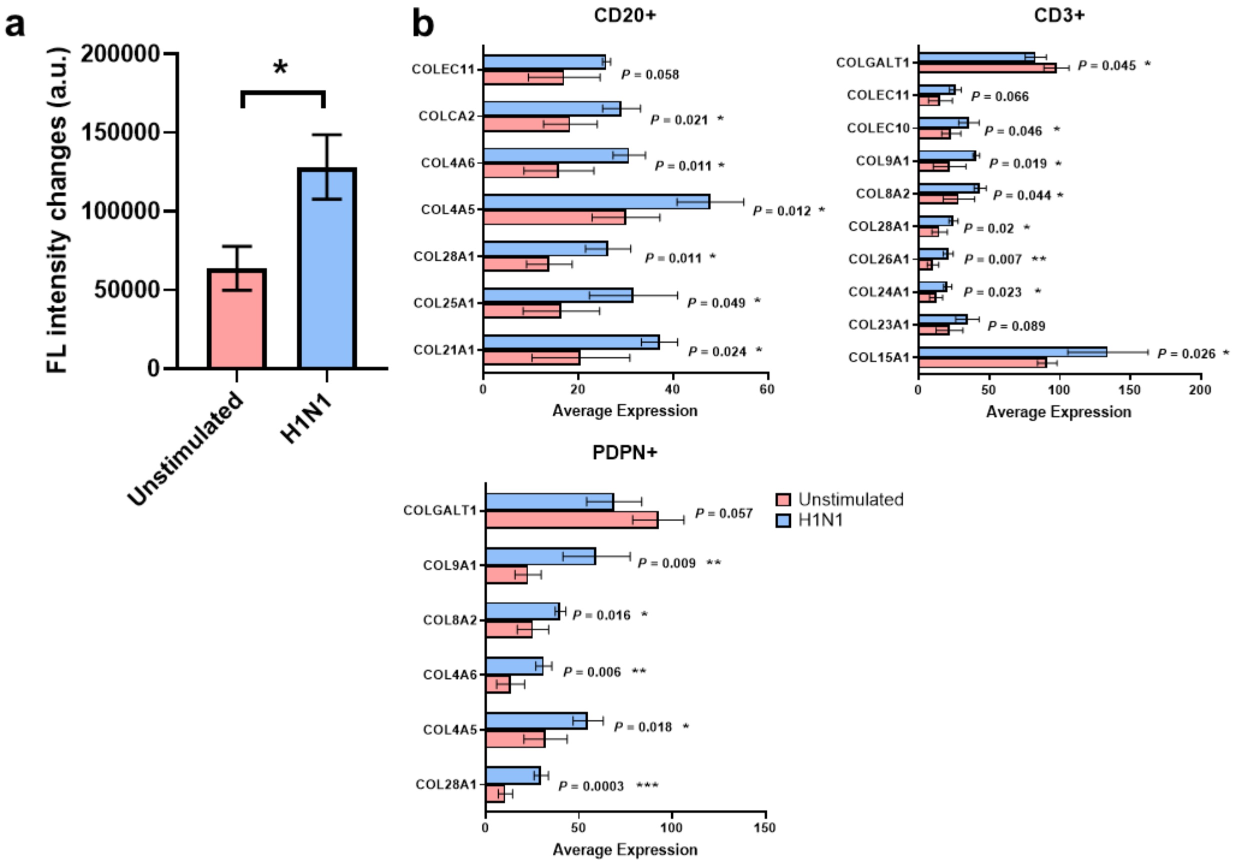

**Figure S7.** Collagen expression in tonsil organoid after H1N1 stimulation. **(a)** Semi-quantification of immunofluorescence images showing the fluorescence intensity changes of COL1 expression in BTO after stimulation with inactivated H1N1. **(b)** Differential gene expression profiles specific to extracellular matrix-related genes in BTO after viral challenge.

**Table S1.** Tonsil donor information

| Donor code | Age (year old) | Gender |
|---|---|---|
| DT008 | 24 | Male |
| DT010 | 29 | Male |
| DT011 | 35 | Female |
| DT013 | 22 | Female |
| DT014 | 37 | Male |
| DT016 | 24 | Male |
| DT018 | 26 | Female |
| DT031 | 33 | Male |

**Table S2.** List of antigens

| Antigen | Source | Working concentration |
|---|---|---|
| Phytohemagglutinin-L (PHA-L) Solution (500X) | Invitrogen | 2.5 µg/ml |
| Lipopolysaccharide (LPS) | Sigma-Aldrich | 50 mg/ml |
| Tetanus toxoid (TT) protein | Creative Biolabs | 0.005 Lf/ml |
| COVID-19 RBD protein | SinoBiological | 2 µg/ml |
| COVID-19 Spike Protein | SinoBiological | 2 µg/ml |
| COVID-19 mRNA vaccine | Pfizer | 0.5 µg/ml |
| SARS-CoV-1 Spike protein | SinoBiological | 10 µg/ml |
| Inactivated Influenza A virus (HA/California/7/2009) | Duke-NUS | 2 µl per 1 ml media |
| R-phycoerythrin (R-PE) | Invitrogen | 2 µg/culture |
| Ovalbumin, Alexa Fluor™ 488 Conjugate | Invitrogen | 4 µg/ml |

**Table S3.** List of reagents used for cytometry by time of flight (CyTOF) analysis

| Reagent | Source | Identifier |
|---|---|---|
| **Antibodies** | | |
| Anti-CCR4, clone 205410 | R & D Systems | Cat# MAB1567 |
| Anti-CCR6, clone G034E3 | BioLegend | Cat# 353402 |
| Anti-CCR7, clone 150503 | R & D Systems | Cat# MAB197 |
| Anti-CD11c, clone B-ly6 | BD Biosciences | Cat# 555391 |
| Anti-CD123, clone 6H6 | Biolegend | Cat# 306002 |
| Anti-CD127, clone A019D5 | BioLegend | Cat# 351302 |
| Anti-CD14, clone M5E2 | BioLegend | Cat# 301802 |
| Anti-CD16, clone 3G8, 209Bi | Standard Biotools | Cat# 3209002B |
| Anti-CD161, clone HP-3G10 | BioLegend | Cat# 339902 |
| Anti-CD185 (CXCR5), clone RF8B2 | BD Biosciences | Cat# 552032 |
| Anti-CD19, clone HIB19 | BioLegend | Cat# 302202 |
| Anti-CD20, clone 2H7 | BioLegend | Cat# 302302 |
| Anti-CD25, clone M-A251 | Biolegend | Cat# 356102 |
| Anti-CD27, clone LG.7F9 | eBioscience | Cat# 14-0271-82 |
| Anti-CD28, clone CD28.2 | BioLegend | Cat# 302902 |
| Anti-CD294 (CRTH2), clone BM16 | BioLegend | Cat# 350102 |
| Anti-CD3, clone UCHT1 | BioLegend | Cat# 300402 |
| Anti-CD38, clone HIT2 | BioLegend | Cat# 303502 |
| Anti-CD4, clone SK3 | BioLegend | Cat# 344602 |
| Anti-CD45, clone HI30, 89Y | Standard Biotools | Cat# 3089003 |
| Anti-CD45RA, clone H100 | BioLegend | Cat# 304102 |
| Anti-CD45RO, clone UCHL1 | Biolegend | Cat# 304202 |
| Anti-CD56, clone NCAM16.2 | BD Biosciences | Cat# 559043 |
| Anti-CD57, clone HNK-1 | BioLegend | Cat# 359602 |
| Anti-CD66b, clone G10F5 | BD Biosciences | Cat# 555723 |
| Anti-CD8, clone SK1 | BioLegend | Cat# 344702 |
| Anti-CXCR3, clone G025H7 | Biolegend | Cat# 353733 |
| Anti-HLA-DR, clone L243 | BioLegend | Cat# 307602 |
| Anti-IgD, clone IA6-2 | Biolegend | Cat# 348235 |
| Anti-PDPN, clone NC-08 | BioLegend | Cat# 337002 |
| Anti-PE, clone PE001 | BioLegend | Cat# 408102 |
| Anti-TCR-γ/δ, clone 5A6.E9, PE | Invitrogen | Cat# MHGD04 |
| **Chemicals** | | |
| Antibody Stabilizer | Candor | Cat# 131050 |
| Iridium (Cell-ID Intercalator-Ir 500 mM) | Standard Biotools | Cat# 201192B |
| Paraformaldehyde 16% | Electron Microscopy Sciences | Cat# 50-980-487 |
| EQ Four Element Calibration Beads | Standard Biotools | Cat# 201078 |
| DN3 Antibody Labeling Kits | Standard Biotools | NA |

**Table S4.** Contributions by authors

| | I.R.S. | D.E.K.C. | H.S.C. | N.S.T. | Z.Z.R.L. | K.S.T. | J.J.H.C. | G.G.K.N. | Y.C.F.S. | G.J.D.S. | C.W.T. | A.T. |
|---|---|---|---|---|---|---|---|---|---|---|---|---|
| Idea conception | MAJOR | | | | | | | | | | | MAJOR |
| Funding acquisition | | | | | | | | | | | | MAJOR |
| Experiment design and methodology | MAJOR | SUPPORT | | | | SUPPORT | | | | | | MAJOR |
| Result analysis | MAJOR | | | | | | | | | | | MAJOR |
| Tonsil tissue provision | | MAJOR | | | | | | | | | | SUPPORT |
| Tissue collection and processing | MAJOR | | | | | | | | | | | SUPPORT |
| Cell isolation and characterization | MAJOR | | | | | | | | | | | SUPPORT |
| Organoid culture and stimulation | MAJOR | | | | | | | | | | | SUPPORT |
| Inactivated virus preparation | | | | | | | | MAJOR | SUPPORT | SUPPORT | | SUPPORT |
| Live virus stimulation (under BSL-3 lab) | SUPPORT | | | | MAJOR | SUPPORT | SUPPORT | | | | | SUPPORT |
| Surrogate virus neutralization tests (sVNTs) assay | MAJOR | | | | | | | | | | MAJOR | SUPPORT |
| Flow cytometry analysis | MAJOR | | | | | | | | | | | SUPPORT |
| cyTOF sample preparation | MAJOR | | | | | | | | | | | SUPPORT |
| cyTOF data analysis | SUPPORT | | MAJOR | SUPPORT | | | | | | | | SUPPORT |
| ELISA assay | MAJOR | | | | | | | | | | | SUPPORT |
| Organoid sectioning and immunofluorescence imaging | MAJOR | | | | | | | | | | | SUPPORT |
| GeoMx sample preparation and data analysis | MAJOR | | | | | | | | | | | SUPPORT |
| Manuscript writing | MAJOR | | | | | | | | | | | MAJOR |
| Manuscript improvement | MAJOR | SUPPORT | SUPPORT | SUPPORT | | SUPPORT | | | SUPPORT | SUPPORT | SUPPORT | MAJOR |
| Entire supervision | | | | | | | | | | | | MAJOR |

Level of contribution: MAJOR, SUPPORT
